# The *Bartonella* autotransporter CFA is a protective antigen and hypervariable target of neutralizing antibodies blocking erythrocyte infection

**DOI:** 10.1101/2021.09.29.462357

**Authors:** Lena K. Siewert, Aleksandr Korotaev, Jaroslaw Sedzicki, Katja Fromm, Daniel D. Pinschewer, Christoph Dehio

## Abstract

Antibodies are key to the clearance of *Bartonella* bacteremia, but the mechanisms and targets of protective antibodies are unknown and bacterial evasion strategies remain elusive. We studied experimental *Bartonella taylorii* infection of mice, its natural host, and investigated protective immune responses. Clearance of bacteremia depended on specific antibodies that interfere with bacterial attachment to erythrocytes. Accordingly, antibodies were effective in the absence of complement and Fc-receptors. Moreover, they formed independently of B-cell hypermutation and isotype class switch. The cloning of neutralizing monoclonal antibodies (mAbs) led to the identification of the bacterial autotransporter CFA as a protective antibody target, and vaccination against CFA protected against *Bartonella* bacteremia. MAb binding mapped to a region of CFA that is hypervariable in both human- and mouse-pathogenic *Bartonella* strains, suggesting mutational antibody evasion. These insights further our understanding of *Bartonella* immunity and immune evasion and elucidate mechanisms driving high *Bartonella* prevalence in the wild.

## Introduction

Neutralizing antibodies protect from infectious agents by impeding or inactivating the pathogen’s essential biological functions. Neutralizing antibodies have been best studied in the context of viral infection, where they typically bind to surface structures, thereby interfering with host cell binding or membrane fusion (Burton, 2002). In case of bacterial infections, secreted toxins are common targets of neutralizing antibodies (Oleksiewicz, et al., 2012). Moreover, neutralizing antibodies have been shown to interfere with various processes and steps in bacterial infection such as bacterial attachment to fibrinogen (Hall, et al., 2003), bacterial growth (Barbour & Bundoc, 2001), agglutination and epithelial cell infection (Menozzi, et al., 1996). For instance, neutralizing antibodies against specific parts of the major outer membrane protein of *Chlamydia trachomatis* prevent host cell attachment and thus antagonize the internalization of this obligate intracellular pathogen (Su, et al., 1990).

Bartonellae are Gram-negative, facultative-intracellular pathogens of mammals that are typically transmitted by blood-sucking arthropods (Harms & Dehio, 2012). The hallmark of infection of these highly host-restricted pathogens in their natural mammalian host is the establishment of a long-lasting intra-erythrocytic bacteremia (Seubert, et al., 2001; Chomel, et al., 2009; Minnick & Battisti, 2009; Harms & Dehio, 2012). *Bartonella* infections are of clinical importance, can be life-threatening and present with a broad spectrum of symptoms that depend on the given species and the immune status of the patient (Maguiña, et al., 2009; Maguiña & Gotuzzo, 2000).

The infection of erythrocytes by *Bartonella* is reported to be a three step-mechanism, consisting of attachment, erythrocyte deformation and invasion (Deng, et al., 2018). Several virulence factors involved in this process have been identified (Mitchell & Minnick, 1995; Coleman & Minnick, 2001; Vayssier-Taussat, et al., 2010) and were found essential for successful colonization of the host (Saenz, et al., 2007; Vayssier-Taussat, et al., 2010).

Previous studies on the clearance of *Bartonella* infection leave important knowledge gaps. Observations in wild animals evidenced multiple sequential bacteremic cycles (Abbott, et al., 1997) and a lack of antibodies in infected animals (Kosoy, et al., 1998; Kosoy, et al., 2004). On the contrary, experimental *Bartonella grahamii* infection of mice showed that antibodies mediate clearance of bacteremia (Koesling, et al., 2001). This goes in hand with high human antibody seroprevalence in endemic regions (Cáceres-Ríos, et al., 1995), detectable antibody responses in infected patients (Spach & Koehler, 1998) and reports on high *Bartonella* strain-specific IgG titers in several experimental animal models as determined by ELISA (Kosoy, et al., 1999; Kabeya, et al., 2006; Kabeya, et al., 2009; Vigil, et al., 2010). Experimentally infected animals failed to develop *Bartonella* bacteremia upon re-exposure to the same strain, which is indicative of protective immunological memory (Kosoy, et al., 1999). Besides antibodies (Koesling, et al., 2001), the cellular immune response (Karem, 2000) and notably T cell help (Marignac, et al., 2010) have been suggested to contribute to the clearance of *Bartonella*.

While the mechanisms underlying antibody protection against *Bartonella* remain essentially unstudied, it has long been speculated that neutralizing antibodies might interfere with the infection of erythrocytes (Karem, 2000; Schülein, et al., 2001; Koesling, et al., 2001). Indeed, antisera raised against the flagellum of *Bartonella bacilliformis* and the IalB protein of *Bartonella birtlesii* were shown to prevent erythrocyte infection by the respective species *in vitro* (Scherer, et al., 1993; Deng, et al., 2016). It remains unknown, however, if these antibody specificities are occurring in the natural infection context.

In this study, we explored the adaptive immune response to *Bartonella* in its natural host, using *B. taylorii* infection of mice. We unravel the requirements for protective antibody formation and their mechanism of action. Moreover, we identify a protective surface antigen in *Bartonella*, the characterization of which across human- and mouse-pathogenic *Bartonella* species offers new insights into bacterial immune evasion with likely implications for *Bartonella* prevalence and iterative infection cycles in the wild.

## Results

### B-cell-dependent clearance of Bartonella bacteremia and passive antibody therapy

Studying the mouse model of *Bartonella* infection, we set out to characterize the adaptive immune response with special emphasis on the role B-cells play in clearing bacteremia. To this end, we used *B. taylorii* IBS296, a strain naturally infecting mice (Harms, et al., 2017). We first compared the course of bacteremia in wild-type (WT) C57BL/6 mice to gene-targeted mice lacking either only B-cells (B-cell ko) or both B- and T-cells (Rag1-/-) (Figure 1A). Intradermal (i.d.) infection of WT mice was followed by a brief abacteremic window of 5-7 days. Bacterial titers in blood then rose to peak at 10-12 days post infection (d.p.i.) with approx. 10^5^ cfu/ml blood. The infection was cleared within 50 d.p.i. and relapses were never observed. This infection kinetics resembled earlier reports from small rodent models of *Bartonella* infection (Koesling, et al., 2001; Okujava, et al., 2014). In contrast, Rag1-/- or B-cell ko mice showed lifelong persistent infection, reaching a plateau of up to 10^7^ cfu/ml blood (Figure 1A). The lack of clearance in Rag1-/- and B-cell knock-out mice confirmed earlier findings on the essential role of B-cells in the clearance of *Bartonella grahamii* in mice (Koesling, et al., 2001). To directly assess the efficacy of antibodies in *Bartonella* control we performed transfer experiments with *B. taylorii*-reactive sera of WT mice (Figure 1B). Immune serum transfer to bacteremic B-cell knock-out mice on day 87 post infection caused bacteremia to subside (Figure 1C, D). Further, we established a prophylactic serum transfer protocol (Figure 1E). WT mice obtained naïve or immune serum on day 3 post infection, thus in the abacteremic window. Recipients of naïve serum developed bacteremia, while the blood of mice given immune serum remained sterile (Figure 1F). Thus, antibodies were effective in both clearing and preventing *Bartonella* bacteremia.

**Figure 1:**
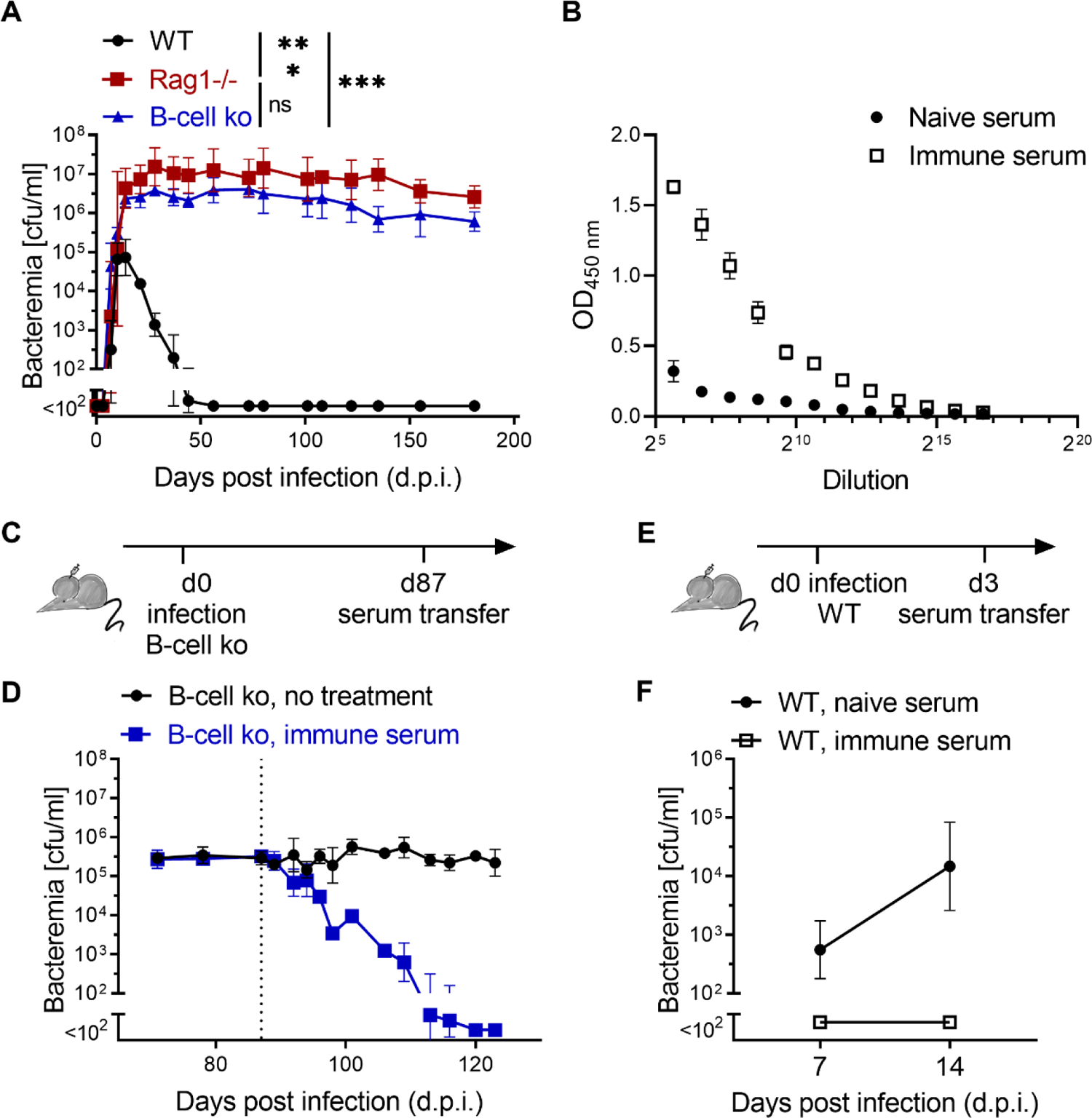
B-cell-dependent clearance of *Bartonella* bacteremia and passive antibody therapy. **(A)** Mice were infected *i.d.* with 10^7^ cfu of *B. taylorii* IBS296 and bacteremia titers (cfu/ml blood) are shown for C57BL/6 wild-type (WT), Rag1-/- mice and B-cell ko mice up to 180 days post infection (d.p.i). **(B)** ELISA assays were performed on plates coated with *B. taylorii* IBS296 outer membrane protein preparation. A pool of immune serum from 10 mice infected with *B. taylorii* mice for at least 45 days was compared to naïve serum. **(C, E)** Schematic of the passive immunization experiments by serum transfer reported in subsequent panels. (C) describes the experiment in (D), (E) describes the experiment in (F). **(D)** We infected B-cell ko mice *i.d.* with 10^7^ cfu *B. taylorii* IBS296 and 87 days later, after establishment of persistent bacteremia, treated them i.v. with 100 µl of either naïve or immune serum raised against *B. taylorii* IBS296 as described in (B). Bacteremia was followed over time; the dashed line indicates the time-point of serum transfer (d87 p.i.). **(F)** WT mice were infected as for the experiment in (D) and were treated i.p. with 100 µl of either naïve or *B. taylorii* IBS296-immune serum on d3. Bacteremia was determined on d14 and d21 p.i.. Data were collected from at least four mice per group, represented as mean ± SD (A, D, F). Data in (B) were collected in technical triplicates. The figures show combined data from two experiments (A, F) or are representative of at least two independent experiments (B, D). Statistical analysis was performed using two-way ANOVA and *P*-values are reported for comparisons evidencing statistically significant differences. Ns: *P* > 0.05; ***: *P* ≤ 0.001

### Immune sera interfere with *Bartonella* adhesion to erythrocytes while complement and Fcγ-receptors are dispensable for bacterial control in mice

We aimed to test the long-standing but so far unproven hypothesis that antibodies interfere with *Bartonella* infection through neutralization defined as the prevention of erythrocyte infection (Karem, 2000; Schülein, et al., 2001; Koesling, et al., 2001). To quantify the protective capacity of immune sera, we modified an *in-vitro Bartonella* erythrocyte infection assay previously published for *B. birtlesii* (Vayssier-Taussat, et al., 2010). Murine erythrocytes were purified and incubated with GFP-expressing *B. taylorii* IBS296 (Figure S1A). Bacterial attachment to erythrocytes was quantified using flow cytometry. To assay for interference of protective antibodies with erythrocyte infection, bacteria were incubated with naïve or immune serum prior to the incubation with erythrocytes (Figure 2A). Immune serum suppressed the rate of GFP+ erythrocytes in a concentration-dependent manner, whereas naïve serum had no observable effect (Figure 2B, C). Heat inactivation of complement in immune serum had no effect on the suppression of erythrocyte adhesion (Figure S1B). No significant bactericidal effect of the immune serum on *B. taylorii* IBS296 was observed by assaying for bacterial growth at the end of the erythrocyte infection assay (Figure S1C), indicating that the reduction of GFP+ erythrocytes by immune serum can unlikely be attributed to killing of the bacteria.

**Figure 2:**
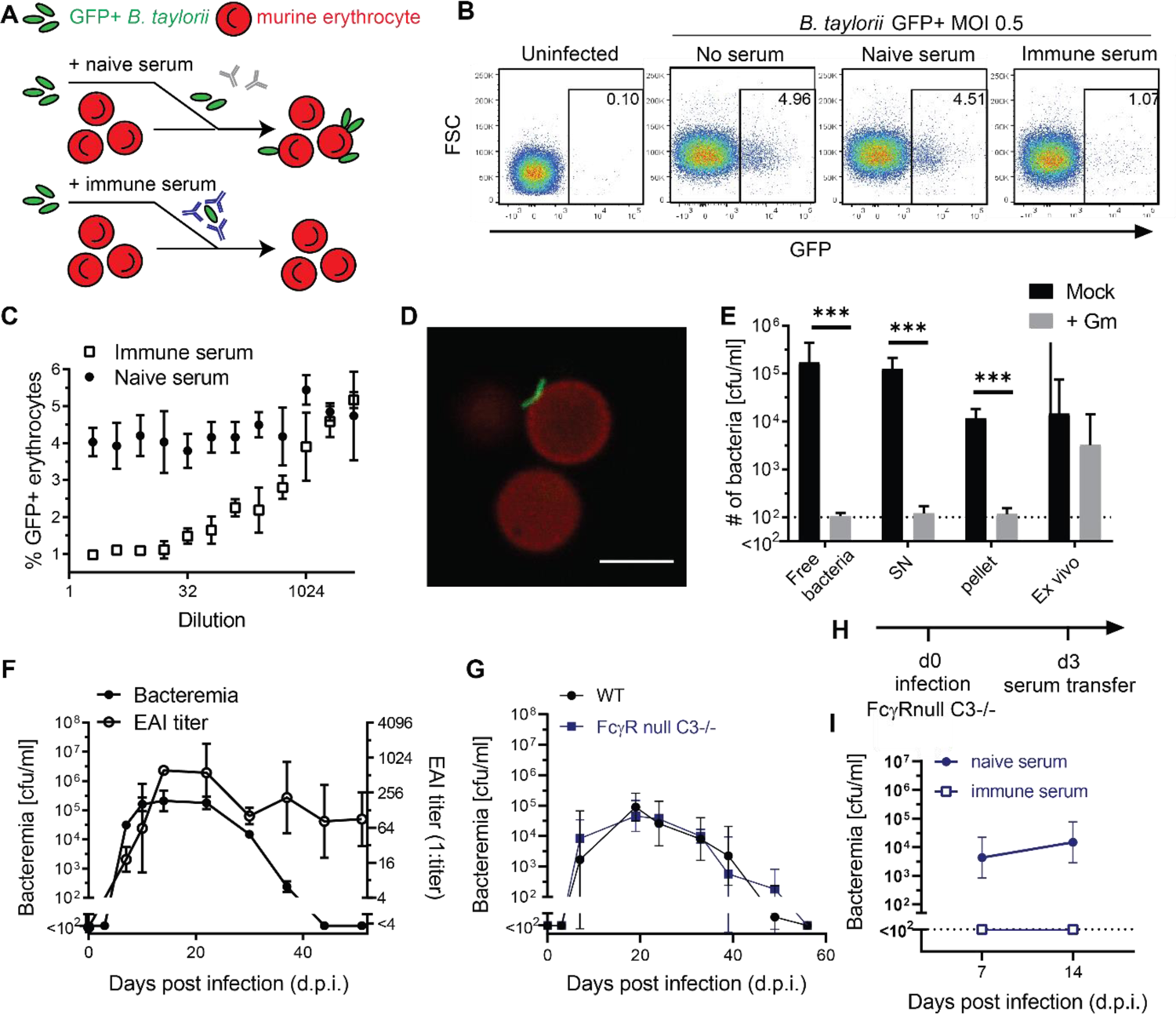
Immune sera interfere with *Bartonella* adhesion to erythrocytes while complement and Fcγ-receptors are dispensable for bacterial control in mice. (**A**) Schematic of the erythrocyte adhesion inhibition (EAI) assay. GFP-expressing *B. taylorii* IBS296 (MOI = 0.5) were incubated with either immune serum or naïve serum for 1 h at 35°C, 5% CO_2,_ subsequently mouse erythrocytes were added. After co-incubation for 24 h, the bacterial adhesion rate to erythrocytes was quantified by flow cytometry. (**B**) Exemplary FACS pseudocolor plots of erythrocytes from the EAI assay: Control erythrocytes without addition of *Bartonella* (uninfected), erythrocytes incubated with GFP-expressing bacteria that have been mock-incubated without serum (no serum) or have been incubated with either naïve (naïve serum) or *Bartonella*-immune serum (immune serum). (**C**) The above EAI assay was conducted using serially diluted serum. The attachment rate in the presence of serially diluted either naïve or immune serum is reported as % GFP+ erythrocytes. (**D**) Confocal images of murine erythrocytes (stained with anti-Ter119 antibody, red) and incubated with GFP-expressing *B. taylorii* IBS296 (MOI = 0.5, green). Scale bar is 5 µm. (**E**) Gentamicin (Gm) protection assays were performed with *B. taylorii* prepared and obtained as follows: Bacteria were either used directly from the plate (free bacteria) or they were incubated with erythrocytes (MOI = 0.5) *in vitro* for 24 h followed by centrifugation to pellet erythrocytes. The resulting supernatant (SN) and pellet were assayed individually. For comparison, erythrocytes were prepared from mice infected with 10^7^ cfu *B. taylorii* IBS296 14 days before (*ex vivo*). (**F**) Bacteremia and EAI antibody titers were monitored longitudinally in WT C57BL/6 infected *i.d.* with 10^7^ cfu *B. taylorii* IBS296. **(G)** Bacteremia titers shown for WT compared to FcγRnull x C3-/- mice. (**H**) Schematic of the serum transfer passive immunization experiments in (I). **(I)** We infected FcγRnull x C3-/- mice *i.d.* with 10^7^ cfu *of B. taylorii* IBS296 and three days later treated them with 100 µl of either *B. taylorii* IBS296-immune (>45 days) or naïve serum. Bacteremia was determined on day 7 and day 14 post infection. Data in (B, C, E) were collected in technical triplicates, one representative FACS plot is shown in (B). (D) Reports one representative image from 50’000 erythrocytes analyzed by fluorescence microscopy. *In vivo* experiments were performed using at least three mice per group. Representative results from one out of two (*in-vivo*) or three (*in-vitro*) similar experiments are shown in (B-G). Pooled data from two independent experiments are shown in (I). Data are reported as mean ± SD. Statistical analysis in (E) was performed using unpaired Student’s *t*-test and statistically significant differences are indicated as ***: *P* ≤ 0.001. See also Figure S1.

Previous studies on *in vitro* infection of erythrocytes showed that *Bartonella* remains extracellular at 24 h post infection (Vayssier-Taussat, et al., 2010). To investigate if our assay is limited to measuring the adhesion to the target cell, we performed confocal microscopy and observed *B. taylorii* IBS296 on the erythrocyte surface only but not in the cells’ interior (Figure 2D). We confirmed this finding using a gentamicin protection assay. Gentamicin does not enter eukaryotic cells and the treatment thus selectively kills extracellular bacteria, while intracellular bacteria survive. Indeed, *B. taylorii* IBS296 remained gentamicin-sensitive during incubation with erythrocytes. In comparison, erythrocytes obtained from infected mice contained mostly gentamicin-insensitive bacteria (Figures 2E and S1E), indicating a mostly intra-erythrocytic localization as previously reported for other *Bartonella* species infecting rodents (Schülein, et al., 2001; Vayssier-Taussat, et al., 2010).

We next aimed to assess the titer of neutralizing serum activity over the course of the infection and tested if the established erythrocyte adhesion inhibition (EAI) assay correlated with clearance. We infected WT mice and followed the animals’ EAI titer over time (Figure 2F). Between day 7 and day 14 post infection, the EAI titer rose sharply and was maintained throughout the bacteremic period and beyond, reaching titers of up to 1:1024. Around the time when EAI serum activity became detectable, the level of bacteria stabilized, and over the subsequent weeks it steadily subsided. The appearance of EAI activity did not, however, result in immediate bacterial clearance. This was in line with the above *in vitro* findings, suggesting protective antibodies interfere with erythrocyte infection prior to bacterial internalization. Bacteria residing inside infected erythrocytes are, therefore, predicted to be protected. Given that the intraerythrocytic stage of *Bartonella* infection is non-haemolytic (Schülein, et al., 2001), a delayed wash-out of intra-erythrocytic bacteremia is compatible with EAI as a main mechanism of anti-*Bartonella* antibody activity.

When following EAI titers after serum transfer into B-cell knock-out mice (Figures 1C, D and S1D), EAI titers reached detection limits within about two to three weeks, and similar kinetics were observed when immune serum was transferred into uninfected B-cell knock-out mice (Figure S1D), suggesting that EAI titers were not substantially masked by antibody binding to free bacteria in circulation. We conclude that antibodies clear the infection by interfering with bacterial attachment to erythrocytes and that the EAI assay can serve for the quantification of *Bartonella*-neutralizing antibody responses.

To test for a potential contribution of antibody Fc-dependent effector mechanisms in bacterial control, we infected mice lacking all α chains of Fcγ-receptors as well as complement component C3 (FcγRnull x C3-/-). In this model, antibodies fail to activate complement or to mediate antibody-mediated cellular phagocytosis and related Fc-dependent effector functions. There was no difference in bacteremia kinetics between FcγRnull x C3-/- and WT mice (Figure 2G), indicating that Fc-dependent effector mechanisms are dispensable and do not significantly contribute to *Bartonella* clearance. Accordingly prophylactic transfer of immune serum protected FcγRnull x C3-/- mice analogously to the observations made in WT mice (Figure 2H, I; compare Figure 1E, F).

### *Bartonella* clearance depends on antibody specificity and CD40L, but occurs independently of antibody affinity maturation and isotype class switching

To further assess the mechanistic requirements for a protective antibody response against *Bartonella* we infected a range of gene-targeted mouse models and measured bacteremia and EAI antibody responses over time. In stark contrast to WT mice, B-cell ko mice failed to mount an antibody response and became persistently infected, as expected (Figure 3A, B; compare Figure 1A, B). B-cells of mice deficient in activation-induced deaminase (AID-/- mice) cannot undergo affinity maturation or class switch recombination, but AID-/- mice nevertheless mounted a largely normal EAI antibody response and controlled *Bartonella* similarly to WT mice (Figure 3C). sIgM-/- mice carry surface IgM on naïve B cells but are unable to secrete soluble IgM and only secrete class-switched immunoglobulin isotypes. Nevertheless, sIgM-/- mice promptly produced *Bartonella*-specific antibodies and controlled the infection (Figure 3D). These data indicated that either IgM or class-switched antibody responses alone sufficed to control *Bartonella* infection, and that the germline antibody repertoire was sufficient to effectively combat this pathogen. AID-/- x sIgM-/- double-deficient mice have B-cells but owing to their combined deficiency are unable to secrete any immunoglobulins. Accordingly, these mice failed to mount antibacterial antibody responses and had unchecked bacteremia for the entire period of observation. While B-cell ko mice are devoid of B-cells as well as antibodies, these findings in AID-/- x sIgM-/- mice documented that it was the antibody production rather than any other role of B-cells in the immune response that controls *Bartonella* infection (Figure 3E). ELISA assays were conducted and confirmed that WT mice produced *Bartonella*-specific IgM and IgG, AID-/- mice produced only IgM whereas sIgM-/- mice were devoid of anti-*Bartonella* IgM but produced class-switched antibodies (Figure S2A). To specifically address a potential role of complement in IgM-mediated bacterial clearance, we tested infection control and antibody responses in AID-/- x C3-/- double-deficient animals but did not find a clear difference in bacterial clearance kinetics or EAI antibody responses (Figure S2B).

**Figure 3:**
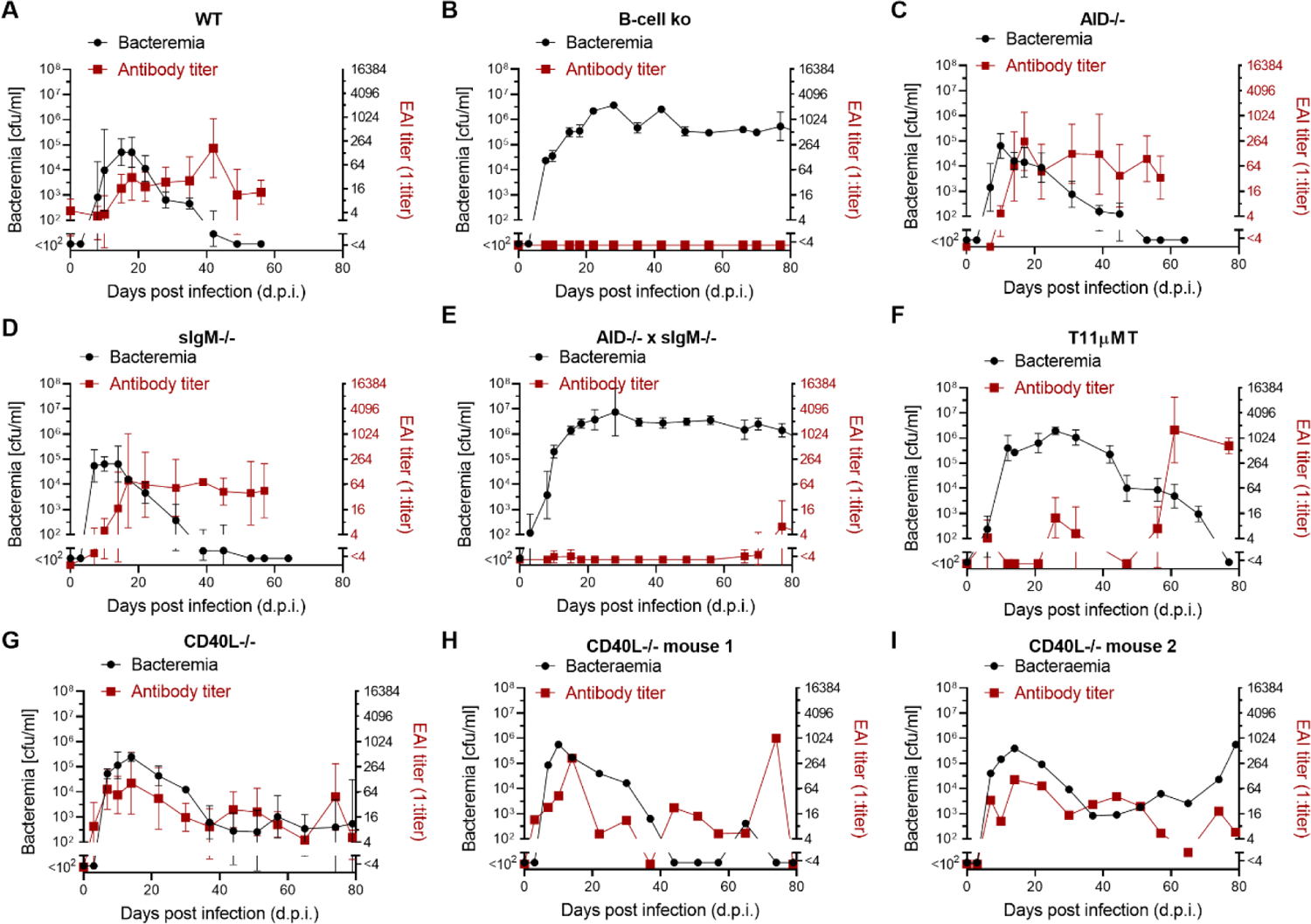
*Bartonella* clearance depends on antibody specificity and CD40L, but occurs independently of antibody affinity maturation and isotype class switching. (**A-G**) We infected groups of mice of the indicated genotypes with 10^7^ cfu *B. taylorii* IBS296 and followed bacteremia as well as EAI titers over time. Panels (**H**, **I**) report these parameters for two individual mice of the experiment shown in (G). Data of additional individual CD40L-/- mice are displayed in Figure S2. Symbols in (A-G) represent the mean of at least four mice per group. Representative data from at least two independent experiments are reported as mean ± SD.

To test whether antibody specificity was important for anti-*Bartonella* defence, we used T11µMT mice, which have an almost monoclonal B-cell repertoire directed against a *Bartonella*-unrelated viral antigen (Bergthaler, et al., 2009). Accordingly, the Bartonella EAI antibody response of T11µMT was substantially delayed. Bacteremia persisted approximately twice as long as in WT mice and only subsided with the advent of a delayed albeit eventually robust antibody response (Figure 3F).

CD4 T-cells have been reported to contribute to *Bartonella* clearance (Marignac, et al., 2010) and T help generally requires T-cell – B-cell interactions involving CD40L – CD40 signalling. In the first thirty to forty days post *Bartonella* infection CD40L-/- mice controlled bacteremia and mounted antibody responses similar to WT mice. At later time points, however, CD40L-/- mice exhibited intermittent drops in antibody titers, which varied between individual mice, and the animals failed to completely eliminate *Bartonella* or evidenced rebound bacteremia after a period of transient bacterial control (Figures 3G-I and S2C-E). Unsustained antibody responses and incomplete pathogen control by CD40L-/- mice mimicked these animals’ behaviour in other infection models (Whitmire, et al., 1996; Bachmann, et al., 2004) and indicated CD40L-dependent helper functions were essential for sustained antibody responses and complete *Bartonella* control.

### Cloning and characterization of protective monoclonal antibodies against *B. taylorii*

In light of this key role of the antibody response in the clearance of *Bartonella* infection, we set out to identify the molecular target(s) of neutralizing antibodies. We generated hybridomas from BALB/c mice immunized with *B. taylorii* IBS296 and identified two promising mAbs that interfered with *Bartonella* adhesion to erythrocytes *in vitro* (LS4G2 and LS5G11; Figure 4A, B). The LS4G2 hybridoma was of the IgG3 isotype but was recombinantly expressed as an IgG2a for comparison to LS5G11. Both antibodies bound to the bacterial surface (Figure 4C, D) yet lacked a detectable bactericidal effect (Figure S3A). Strikingly, both antibodies were highly specific for the *B. taylorii* IBS296 strain used for immunization and failed to bind another *B. taylorii* isolate (strain M1) or other, closely related *Bartonella* species (Figure S3B). Since both antibodies were selected for their *in vitro* activity, we wanted to test if they are also protective *in vivo*. To this end we infected mice with *B. taylorii* IBS296 and 3 days later treated them with either LS4G2, LS5G11 or isotype control antibody. The blood of recipients of LS4G2 and LS5G11 remained sterile, while those given isotype control antibody became bacteremic (Figure 4E, F). These findings indicated that the EAI assay identified antibodies that exhibited *in vivo* protective capacity. Analogous results on LS4G2 passive immunization, were obtained in FcγRnull x C3-/- mice (Figure 4G; H), further supporting the conclusion that LS4G2 protected by neutralizing *Bartonella* independently of classical complement or Fc-mediated effector functions.

**Figure 4:**
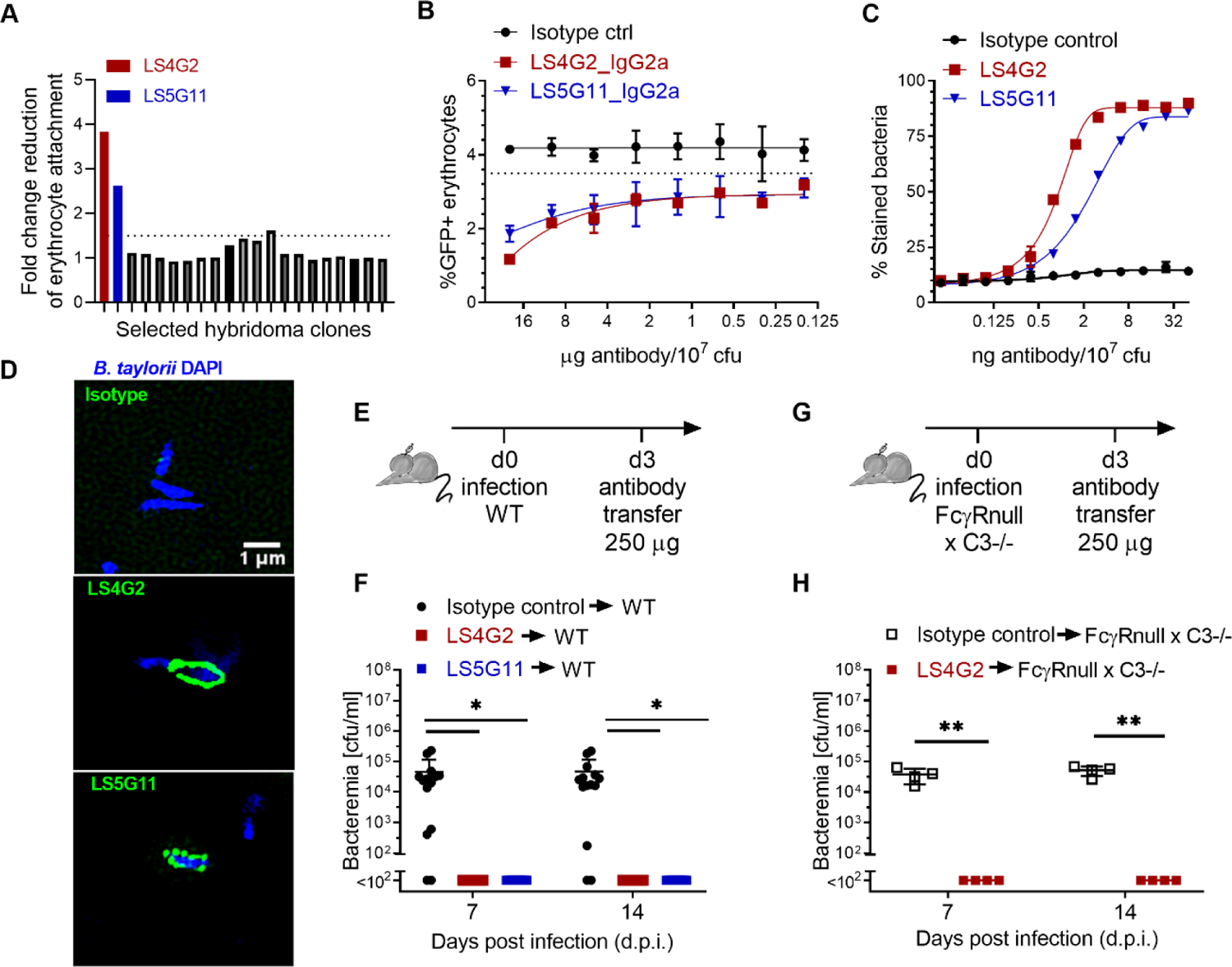
Cloning and characterization of protective monoclonal antibodies against *B. taylorii* IBS296. (**A**) Supernatants collected from 20 individual hybridoma clones in our screen were analyzed using EAI assays. The potency of each clone is expressed as fold-change reduction of erythrocyte attachment when compared to mock (medium only). The two most potent clones (LS4G2 and LS5G11) are highlighted in red and blue, respectively. (**B**) The EAI potency of purified LS4G2 (red) and LS5G11 (blue) in side-by-side comparison to isotype control antibody. (**C**) *B. taylorii* IBS296 surface staining by LS4G2 (red), LS5G11 (blue) and isotype control antibody was quantified by flow cytometry. (**D**) Surface labelling of *B. taylorii* IBS296 (DAPI, blue) upon incubation with LS4G2, LS5G11 or isotype control antibody (all in green) was visualized by structured illumination microscopy. (**E, G**) Schematic of passive immunization antibody transfer experiments conducted in panel F (schematic in E) and panel H (schematic in G): WT (F) and FcγR null x C3-/- mice (H) were infected *i.d.* with 10^7^ cfu *B. taylorii* IBS296 and three days later were treated with 250 µg of the indicated antibodies *i.v..* Bacteremia was determined on day 7 and on day 14 post infection. (B-D) One representative data set from two independent experiments is shown. Each experiment was performed in technical triplicates. (F) Pooled data from three independent experiments are reported. Symbols in (F, H) represent individual mice (n=10 per group in F, n=4 per group in H). Statistical analysis was performed by unpaired Student’s *t*-tests, *: *P* ≤ 0.05, **:*P*≤ 0.01. Error bars show SD. See Fig. S3 for related data.

### The *B. taylorii* autotransporter CFA is a target of protective antibodies

To identify the bacterial targets of the two protective mAbs, we performed immunoprecipitations from the solubilized outer membrane fraction of *B. taylorii* IBS296 followed by mass spectrometry. Surprisingly, both antibodies pulled down the same protein: OPB34894.1 (Figure 5A, B). The orthologue in *B. henselae* was previously identified as an autotransporter with a putative co-hemolysin activity and has been termed CAMP-like factor autotransporter (CFA) (Litwin & Johnson, 2005). Autotransporter proteins constitute a large family of virulence factors secreted by Gram-negative bacteria via the type V secretion mechanism. They are characterized by an extracellular passenger domain at the N-terminus that often remains anchored in the outer membrane via a C-terminal β-barrel (Tame, 2011). Deletion of the *cfa* locus in *B. taylorii* IBS296 resulted in complete loss of binding by LS4G2 and LS5G11, a phenotype which could be rescued by the expression of the protein from a plasmid (Figure 5C). Finally, the expression of CFA of *B. taylorii* IBS296 (CFA_Btay_) in other *Bartonella* species, which in their native form failed to react with our mAbs, resulted in surface staining (Figure S3B-D). Altogether, these observations validated CFA as the molecular target of the protective antibodies LS4G2 and LS5G11.

**Figure 5:**
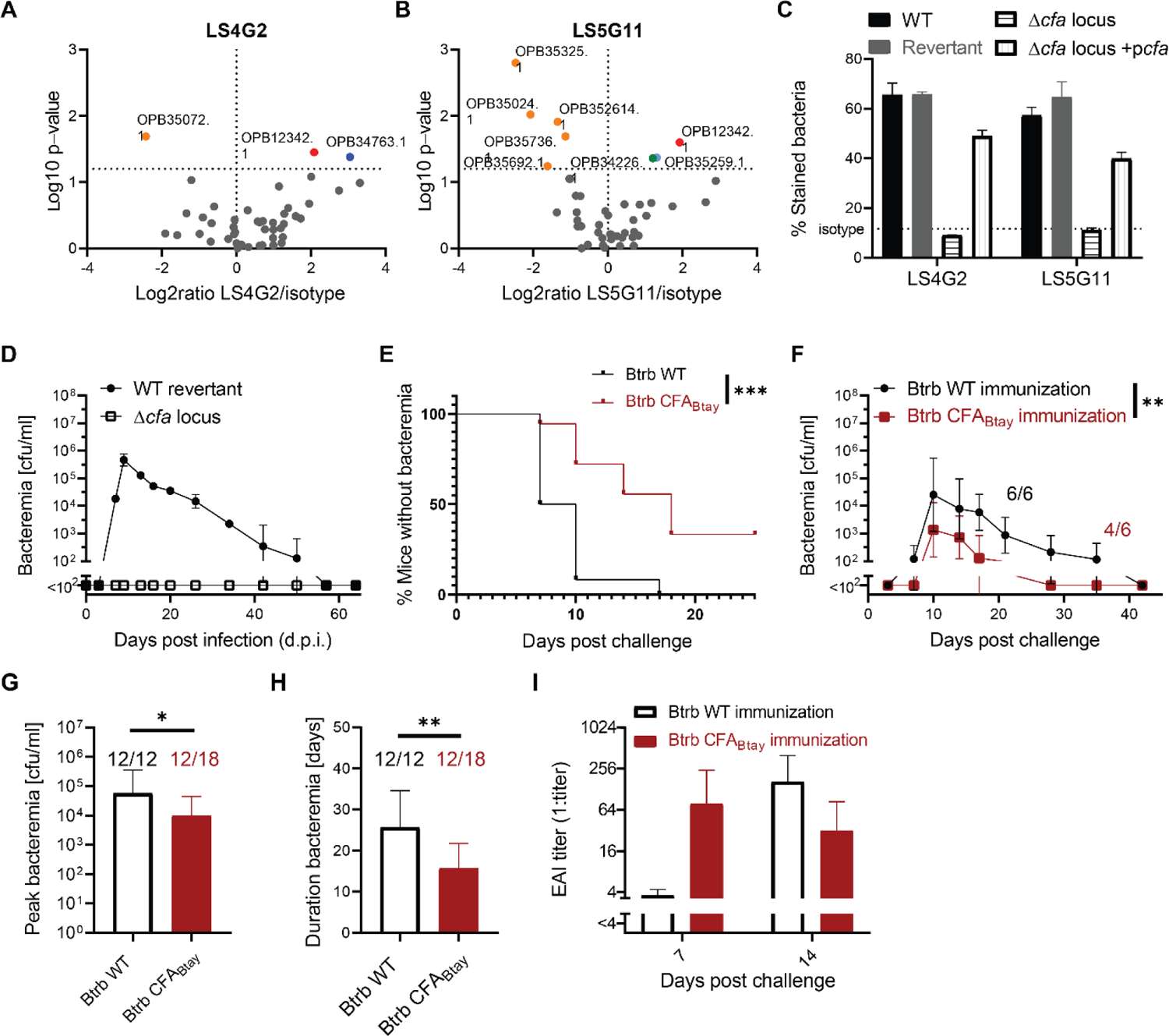
The *B. taylorii* autotransporter CFA is a target of protective antibodies. (**A**, **B**) Volcano plots of proteins identified by mass spectrometry after pull-down with either LS4G2 (A) or LS5G11 (B). OPB12342 denominates the gene product of *cfa*. The second LS4G2 hit OPB34763.1 could not be confirmed in a repeat experiment and hence was not pursued further. (**C**) To validate CFA as molecular target of the LS4G2 and LS5G11 mAbs we stained the following versions of *B. taylorii* IBS296 and determined surface-bound antibody by flow cytometry: i) the parental WT bacterium, ii) the *cfa* locus knock out mutant (Δc*fa* locus), iii) the corresponding WT “revertant” strain, and iv) the *cfa* locus knock out strain complemented by expression of *cfa* from plasmid (Δc*fa* locus + p*cfa*). Note that the WT revertant and the Δc*fa* locus knock mutant are isogenic strains, which are progeny of the same bacterial mutagenesis intermediate but represent the result of recombination events in different homology regions, which both led to deletion of a transiently integrated mutagenesis plasmid. Symbols represent the mean ± SD of 3 technical replicates. (**D**) We infected WT C57BL/6 mice (n=3 per group) *i.d.* with 10^7^ cfu of either WT revertant or Δc*fa* locus deletion mutant of *B. taylorii* IBS296 and monitored bacteremia over time. (**E-I**) We immunized C57BL/6 WT mice *i.d.* either with 10^7^ cfu of WT *B. tribocorum* (Btrb WT) or with the same dose of an isogenic strain expressing CFA of *B. taylorii* IBS296 (Btrb CFA_Btay_). Six weeks later both groups of mice were challenged with *B. taylorii* IBS296. (**E**) The percentage of abacteremic mice as a function of time. Combined data from 3 experiments with 3-6 mice per group each. (**F**) Bacterial loads in blood are selectively reported only for bacteremic animals (6/6 Btrb WT-immunized mice; 4/6 Btrb CFA_Btay_-immunized mice) from one out of three experiments. (**G, H**) Peak bacteremia (G) and duration of bacteremia (H) for bacteremic animals (12/12 Btrb WT-immunized mice, 12/18 Btrb CFA_Btay_-immunized mice). Combined data from three independent experiments. **(I)** EAI titers of the animals in (F) were determined on day 7 and on day 14 after *B. taylorii* challenge. (A-D) Representative data from two independent experiments are shown. For statistical analysis we used the Log-rank test (E), two-way ANOVA (F) and unpaired Student’s t-test (G H). *: P ≤ 0.05; **: P≤ 0.01; ***: P ≤ 0.001. Data in (C, D, F, I) are reported as mean ± SD, mean ± SEM is shown in (F).

CFA has been shown essential for *B. tribocorum* bacteremia in rats (Saenz, et al., 2007) and *B. birtlesii* bacteremia in mice (Vayssier-Taussat, et al., 2010). We extended these findings to *B. taylorii* IBS296 by infecting mice with either one of two isogenic strains, namely the deletion mutant and its revertant wild-type variant. Indeed, the blood of the animals infected with the Δ*cfa* locus mutant remained sterile, while revertant wild-type bacteria caused normal bacteremia (Figure 5D).

*B. tribocorum* naturally infects rats but does not cause robust bacteremia in mice (Schülein, et al., 2001). Here we used it as a prototypic live vaccine vector to test whether CFA is a protective antigen in *B. taylorii* IBS296. We vaccinated mice with an isogenic *B. tribocorum* strain, which we engineered to ectopically express CFA_Btay_, whereas control animals were administered WT *B. tribocorum*. Six weeks post immunization we challenged both groups of mice with *B. taylorii* IBS296.

All control mice immunized with WT *B. tribocorum* developed *B. taylorii* IBS296 bacteremia. In contrast, one-third of the mice vaccinated with *B. tribocorum* expressing CFA_Btay_ remained abacteremic (Figure 5E). The other two-third of vaccinated mice, albeit developing bacteremia, exhibited a significantly reduced peak bacterial burden and a shortened duration of bacteremia (Figure 5F-H). Importantly, mice immunized with *B. tribocorum* expressing CFA_Btay_ developed EAI antibodies within the first 7 days after challenge, whereas the control group only did so by day 14 (Figure 5I). Taken together, these experiments identified the autotransporter CFA as a protective antibody target and candidate vaccine antigen in *Bartonella*.

### High variation of CFA on the sub-species level suggests immune evasion from antibody selection pressure

We next performed a comparative genomics analysis among the Bartonellae to learn more about the structure of the *cfa* locus as well as about the level of conservation and sequence variation in CFA. We found that the *cfa* locus is present in all Eubartonellae (Figure S3E). The locus is composed of the *cfa* gene, immediately followed by a variable number of downstream genes or pseudogenes with homology to *cfa*, suggesting that those represent remnants of gene duplication or recombination events. The *cfa* locus is flanked by synteny blocks consisting of highly conserved genes responsible for iron uptake (*fatBCD*, located upstream) and accountable for creatin and cobalamin metabolism (BH12990-12970, located downstream), respectively (Figure 6A).

**Figure 6:**
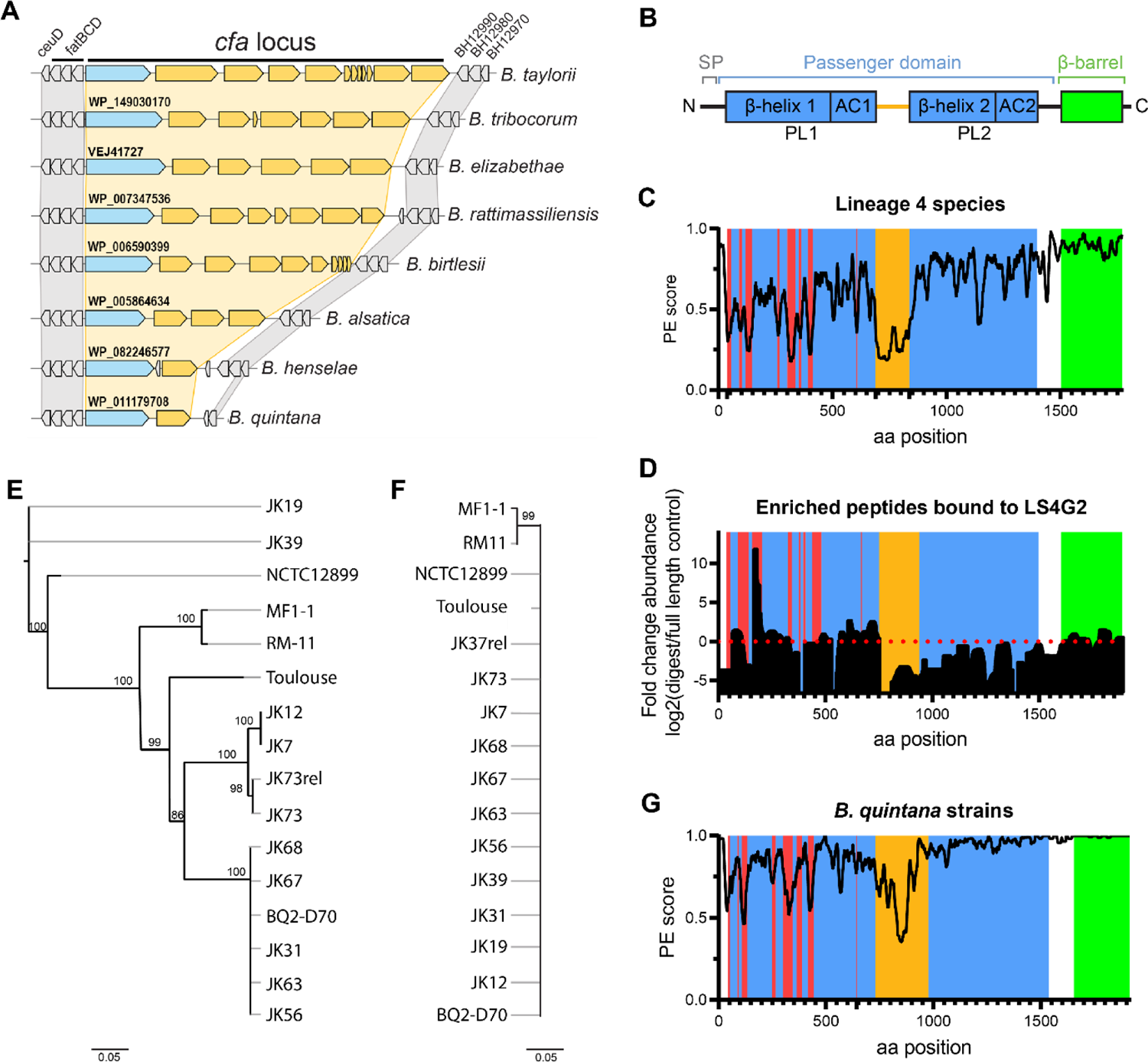
High variation of CFA on the *Bartonella* sub-species level suggests immune evasion from antibody selection pressure. (**A**) Gene synteny analysis of the *cfa* locus and flanking regions in the indicated *Bartonella* species. Flanking synthenic housekeeping genes are shown in grey, the *cfa* locus in blue with the corresponding gene annotation and further putative autotransporter genes/pseudogenes in yellow. (**B**) Domain organization of CFA of *B. taylorii* IBS296. SP (signal peptide, grey), the surface-located passenger domain with two β-helices (blue), pertactin-like domains (PL) and corresponding autochaperones (AC) predicted to form two stalks, separated by a linker (orange) and the C-terminal β-barrel (green). (**C**, **G**) Conservation scores of amino acid sequences along alignments of CFA homologs for (**C**) representative species of *Bartonella* lineage 4 or (**G**) closely related strains of *B. quintana*. PE (Property Entropy) score is a metrics based on Shannon Entropy (Capra & Singh, 2007). Colouring scheme is based on colours used to illustrate CFA domain organization in (B), except for red colour indicating hypervariable regions. (**D**) Analysis of putative LS4G2 binding regions in CFA of *B. taylorii* IBS296. After LS4G2 immunoprecipitation of CFA from bacterial lysates, a partial digest with proteinase K was performed and the eluted peptides were compared to full length eluted (undigested) protein by mass spectrometry. Data are represented as fold change difference between the above two conditions. The domain architecture is depicted including regions of high variability between *B. taylorii* strains IBS296 and M1 (red). Representative data from two independent experiments are reported, each of them performed in biological triplicates. (**E**, **F**) Phylogeny analysis of the indicated *B. quintana* strains were generated for the genes encoding either *cfa* (**E**) or *virB5* (**F**). Values above nodes show bootstrap support (>75%). Scale bar indicates number of substitutions per site. See Figures S4 and S5 for further detail.

Protein folding predictions indicated that the passenger domain of CFA is composed of two β-helices separated by a disordered linker (Figure 6B). Each β-helix contains a predicted pertactin-like (PL1, PL2) fold including a C-terminal autochaperone (AC1, AC2) (Rojas-Lopez, et al., 2018).

Comparative sequence analysis of CFA across lineage 4 species indicated that the level of sequence conservation of the passenger domain is inhomogeneous. We found that the N-terminal stalk, which is likely the distal part of CFA exposed on the bacterial surface, contains eight variation hot spots (V1-V8; red in Figures 6C, S4 and S5A) and thereby overall is less conserved than the **C-**terminal stalk located closer to the membrane anchor. Structure prediction indicated that these variable regions overlap with parts of the polypeptide chain that do not form structural elements of the β-helix fold but rather decorate its surface, likely projecting outwards (Figure S5B, C).

We next aimed at mapping the LS4G2 epitope on CFA. We performed immunoprecipitation of CFA followed by limited proteolysis with proteinase K and compared the eluted peptides to the undigested full-length protein by mass spectrometry (Figure 6D). This approach revealed a supposed LS4G2 binding region in the distal (N-terminal) half of the passenger domain, located around amino acid positions 170 to 190. This region overlaps with one of the variation hotspots (V3) (Figure S4A, S5C), which exhibits substantial sequence differences between *B. taylorii* isolates IBS296 and the M1 isolate that is not bound by LS4G2 (Figure S3B, S4A, S5C) and which we identified in our comparative analysis of lineage 4 species (Figure 6C, highlighted in red; Figure S4B). This finding suggested that variation in this antigenic region of CFA may facilitate immune escape. We aimed to generalize and extend these findings to the human pathogen *B. quintana* (Figure 6E-G, Figure S4C), for which a large number of closely related isolates have been collected from patients in San Francisco during the AIDS epidemic (Table S1). We found that the pattern of variability hotspots on the sub-species level was similar to that observed at the species level in lineage 4 (Figures 6G, compare Figure 6C; Figures S4B, C). Strikingly, the variability of CFA sequences was significantly higher than the one in VirB5, an important virulence factor of *Bartonella*, which remained almost unchanged at the strain level (Figure 6E, F). Collectively, our findings document high variability in the surface-exposed regions of CFA and demonstrate these same regions might be the targets of neutralizing antibodies. Taken together, this pinpoints to antibody selection pressure accelerating the evolution of this protein domain, representing supposed hotspot of mutational immune escape.

## Discussion

Antibodies represent a central element of adaptive immunity against many microbial invaders. The same applies obviously to *Bartonella* infection, yet the kinetics of protective antibody responses, their mechanism of action and molecular targets but also the immunological processes underlying the formation of *Bartonella*-specific antibody immunity have long represented an understudied subject. By addressing these fundamental knowledge gaps, the present work redefines our general understanding of *Bartonella* – host interactions, sharpens concepts of the germ’s habitat in mammalian hosts and offers prospects on vaccine design.

We suppose CFA is a key molecular target of protective antibodies, although perhaps not the only one. Its sequence hypervariability in antibody binding sites indicates that *Bartonella* is under antibody selection pressure in the wild and evades it at the sub-species level of individual strains. We propose that CFA hypervariability may facilitate multiple sequential or even timewise overlapping infection episodes of the same mammalian host by closely related *Bartonella* strains. Besides enlargement of the bacterial habitat in time and host space, superinfection as a consequence of antibody evasion should create opportunities for gene transfer between different *Bartonella* strains as discussed below and thus is expected to expedite the evolution of the species.

From a vaccinology standpoint, the hypervariability of a key protective antibody target heralds major challenges. CFA hypervariability resembles the immune evasion strategy of microbes such as the malaria parasite *Plasmodium falciparum* with its vast antigenic variation in adhesion molecules (Su, et al., 1995; Gardner, et al., 2002), the extensive diversity in streptococcal cell wall polysaccharides (Wu, et al., 2015) or gonococcal pilus variants (Meyer & van Putten, 1989), but also the far over one-hundred known rhinovirus serotypes resulting from viral capsid protein variation (Lewis-Rogers, et al., 2014). Unlike for influenza A virus, these pathogens do not depend on their serial replacement over time by new antigenic variants. On the contrary, a large number of variants co-circulate in host communities, raising the bar for a protective vaccine. We report that immunization with vectored CFA matching the *Bartonella* challenge isolate affords protection against bacteremia.

For clinical use in humans, however, a vaccine would ideally cover all important pathogenic species, or should at least encompass most strains of a given high-incidence species, such as *B. henselae* or *B. quintana* that are responsible for most human infections worldwide. Vaccination may thus have to focus antibody responses on conserved regions and epitopes of CFA, provided such antibodies can be elicited, resembling ongoing efforts in epitope-focused vaccine design for HIV, hepatitis C virus (HCV) and influenza A virus (Correia, et al., 2014).

It remains to be investigated whether *Bartonella* carries invariant or less variable protective targets than CFA, which can more easily be exploited for vaccine design. Both protective antibodies identified in our study target CFA, but in light of the limited number of antibodies screened it remains entirely possible that *Bartonella* exhibits additional protective antibody targets. Besides the numeric limitations of our screen, the screening method relied on bacteria grown on agar plates, such that alternative antibody targets would have escaped detection if their expression was restricted to infection conditions. Similarly, antibodies interfering with steps of the bacterial life cycle other than erythrocyte adhesion would have gone undetected in our EAI assay.

CFA is present in all Eubartonellae yet its variability extends to the level of individual strains, indicating immune escape rather than host adaptation drives its evolution. The genomic organization of the *cfa* locus, with the *cfa* gene followed by multiple homologous genes, gene fragments and/or pseudogenes, suggests that recombination rather than the accumulation of single base pair changes is driving this process. The lack of recurrence of bacteremia in the experimental model suggests further that immune evasion depends on horizontal gene transfer between a larger pool of variant sequences that have evolved in closely related strains coexisting in the wild. The gene transfer agent (GTA), conserved in all Bartonellae, mediates high frequency horizontal gene transfer (Québatte, et al., 2017; Québatte, 2019) and might contribute to antigenic variation of CFA with resulting immune escape. CFA and other autotransporter proteins were found essential for colonization of the host (Vayssier-Taussat, et al., 2010; Saenz, et al., 2007). The present findings suggest protective CFA-binding antibodies capture bacteria when seeded into the bloodstream and prevent their attachment to erythrocytes, a key step in the bacterial life cycle. Future studies should address how Bartonellae diversify the antibody binding regions of CFA at high frequency, while safeguarding its functionality for the infection process.

Taken together this study provides fundamental conceptual insights into *Bartonella* immune control and immune evasion in rodent hosts. Our findings have far-reaching implications for the understanding of the bacterial niche in its natural habitat and delineate challenges in vaccine design related to newly identified antigenic variation at the *Bartonella* species level.

## Acknowledgements

We thank Alexander Harms, Marianna Florova and Alexander Wagner for their valuable input into planning the experimental design and approaches. We thank Stefan Bienert who made the AlphaFold 2 pipeline available on our local infrastructure in sciCORE and we are grateful to Richard Neher for helping to run basecalling and barcoding in Guppy software. This work was supported by the Swiss National Science Foundation (SNSF, www.snf.ch) grants No. 310030B_201273 to C. D. and 310030_173132 to D. D. P., the National Centre of Competence in Research (NCCR) AntiResist funded by SNSF grant number 51NF40_180541 to C. D., as well as by the European Research Council (ERC) grant No. 310962 to D. D. P..

## Author Contributions

L. K. S, D. D. P. and C. D. conceptualized all studies and designed all experiments. L. K. S. (animal work, EAI assay, flow cytometry, mass spectrometry, hybridoma generation and antibody production, microscopy, cloning); A. K. (bioinformatic analysis); J. S. (cloning, microscopy) and K. F. (animal work) conducted the experiments, collected and analyzed the data. L. K. S., J. S., D. D. P. and C. D. wrote the manuscript. All authors have read and approved the final version of the manuscript.

## Material and Methods

### Bacterial strains and growth conditions

All bacterial strains and plasmids used in this study can be found in the key resource table and supplementary table S3. *E. coli* strains were cultivated in lysogeny broth (LB) or solid agar plates supplemented with the appropriate antibiotics at 37°C overnight.

*Bartonella* strains were grown at 35°C and 5% CO_2_ on Columbia blood agar (CBA) plates supplemented with 5% defibrinated sheep blood and the appropriate antibiotics. *Bartonella* stocks are maintained at −80°C and are streaked as “thumbnails” for 3 days and are subsequently expanded for 2 days prior to the experiment.

Plasmids were introduced into *Bartonella* strains by conjugation from *E. coli* strain β2150 using biparental mating (Harms, et al., 2017).

Antibiotics or supplements were used in the following concentrations: kanamycin at 30 µg/ml, gentamicin at 10 µg/ml, streptomycin at 100 µg/ml, diaminopimelic acid (DAP) at 1 mM.

### Mouse strains and husbandry

Female BALB/cJRj and C57BL/6JRj mice were purchased from Javier Labs.

The genetically modified strains AID-/- (Muramatsu, et al., 2000), β2M-/- (Zijlstra, et al., 1989), C3-/- (Wessels, et al., 1995), JHT-/- (Chen, et al., 1993), K*^b^*D*^b^*-/- (Vugmeyster, et al., 1998), MHCII-/- (Madsen, et al., 1999), Rag1-/- (Mombaerts, et al., 1992), sIgM-/- (Tsiantoulas, et al., 2017), TCRβδ-/- (Mombaerts, et al., 1993) and T11µMT (Klein, et al., 1997) were bread at the Laboratory Animal Science Center (LASC, University of Zurich, Switzerland) under SPF conditions. FcγR4alpha x C3 -/- (crossing of C3-/- with FcγR4alpha-/- (Fransen, et al., 2018)) were bread at the Transgenic Mouse Core Facility (TMCF, University of Basel, Switzerland). µMT-/- mice (Kitamura, et al., 1991) were obtained from Jackson Laboratories, Maine, USA. The strains CD1d-/- (Smiley, et al., 1997) and MR1-/- (Treiner, et al., 2003) were a kind gift from Prof. Genaro De Libero, University of Basel. AID x C3-/- was obtained by crossing AID-/- and C3-/-. TCRβ-/- and TCRδ-/- single knock outs were obtained by back-crossing TCRβδ-/- with WT C57/BL6. CD1d-/- x MHCII-/- and MR1-/- x MCHII-/- were obtained by crossing the respective line with MHCII-/- mice.

All mice were in C57BL/6 background unless specified otherwise. Adult mice (5-8 weeks) of both sexes were used for experiments.

Mice were housed in ventilated cages on sterilized bedding and provided water and food *ad libitum*. Housing density was maximal five mice per cage and mice were allowed to acclimatize for at least a week undisturbed. The animal room was on a 12 light/ 12 dark cycle, and cage bedding changed every week. Mice were housed in strict accordance with the Federal Veterinary Office of Switzerland and/or local animal welfare bodies. All animal work was approved by the Veterinary Office of the Canton Basel City (license no. 1741).

## Method Details

### Constructions of strains and plasmids

DNA manipulations were performed according to standard techniques and all cloned inserts were DNA sequenced to confirm sequence integrity.

For the generation of bacterial knock outs the previously described two-step selection procedure for gene replacement was used (Schulein & Dehio, 2002). For complementation overexpression selected genes from plasmid were cloned into variants of the plasmids pJS43 (Celli, et al., 2005) under the control of the *AphT* promotor of pJC43 (GFP-expression) or IPTG-inducible promotor P*lac*(MQ5) (Harms, et al., 2017) (CFA_Btay_). For genetic complementation under the natural promotor in the chromosome insertions via the Tn7 transposon were used (based on pUC18T-mini- Tn7, (Choi, et al., 2005)).

For antibody expression vectors, the corresponding V regions of heavy and light chain were cloned into the CMV-promoter-driven mammalian expression vector pXLG1.2, followed by the Cγ2a constant domain (kindly provided by Prof. Shozo Izui, University of Geneva).

A detailed description for the construction of each plasmid is presented in Table S3. The sequences of all oligonucleotide primers and synthesized gene fragments used in this study are listed in Table S2 and S4.

### Animal experimentation

Animals were infected or immunized *i.d.* with 10^7^ cfu bacteria in PBS under isofluorane/oxygen anaesthesia. Blood was drawn in 3.8% sodium citrate at the indicated days post infection. For blood cfu count, whole blood was frozen at −80°C, thawed and plated in limited dilution series on CBA blood agar. For serum analysis, the blood samples were centrifuged for 5 min at RT at 5000 x g in serum tubes. Serum was frozen at −20°C until further usage. Immune serum was obtained 45 dpi from immunized C57BL/6 mice and pooled from 20 animals. Naïve or immune serum were injected *i.v.* in 100 µl. Antibodies were given *i.v.*in a dose of 250 µg in PBS.

### Erythrocyte infection and EAI assay

Murine Erythrocytes were isolated from BALB/c mice after blood collection in 3.8% sodium citrate by terminal bleeding. Erythrocytes were purified using a Ficoll-gradient and kept at 4°C in DMEM supplemented with 10% FCS for up to two weeks until usage.

For the EAI assay, a serial dilution of sera or antibodies was performed in a U-bottom 96-well plate in DMEM with 10% FCS. 5*10^5^ bacteria (GFP+) were added per well and incubated for 1 h at 35^°^C and 5 % CO_2_. 10^6^ (MOI 0.5) erythrocytes were added in 100 µl DMEM containing 10% FCS. After 24 h the supernatant was removed and the cells were fixed in 1% PFA and 0.2% GA in PBS for 10 min at 4°C. FACS buffer (2% FCS in PBS) was added and the plates were analyzed by flow cytometry (CantoII, using the HTS autosampler). If indicated, the serum was heat-inactivated at 60°C for 30 min before usage.

EAI titers were calculated by endpoint titer determination as described (Frey, et al., 1998).

For confocal microscopy, erythrocytes were infected using the same conditions lacking the serum pre-incubation.

### Microscopy

Bacteria were grown as described above, collected in PBS and stained with purified antibody for 45 min at RT. Bacteria were then washed twice with PBS, centrifuged at 4000 x g for 5 min and stained with the secondary antibody (anti-mouse IgG-Alexa 488) for 1 h at 4°C in the dark. After 3 washes, the bacteria were stained with DAPI for 1 h at 4°C in the dark and after another three washes fixed in 3.7% PFA for 10 min at 4°C in the dark.

Erythrocytes were infected with GFP+ bacteria as described above. Erythrocytes were then stained with anti-Ter119-Alexa 647 for 1 h at 4°C in the dark. After washing and centrifugation at 100 x g, 5 min, the cells were fixed in 1% PFA and 0.2% GA in PBS for 10 min at 4°C in the dark.

Fixed erythrocyte samples were centrifuged at 100 x g for 5 min. Stained bacteria samples were centrifuged at 4000 x g for 5 min. For both types of samples, the supernatant was removed and the cell pellet was resuspended in ProLong Diamond Antifade mountant. 15 µl of the suspension was applied on 18 mm, #1.5 thickness coverslips and mounted onto glass slides. After 24 h of curing at room temperature, the coverslips were sealed with nail polish.

Confocal images were acquired with a SP8 confocal microscope (Leica) equipped with 488 and 638 nm solid-state lasers and a Plan Apo CS2 63x, 1.40 NA oil objective.

3D-SIM was performed using a DeltaVision OMX-Blaze system (version 4; GE Healthcare) equipped with 488 and 568 nm solid-state lasers, Plan Apo N 63x, 1.42 NA oil objective and 4 liquid-cooled sCMOs cameras (pco Edge, full frame 2560 x 2160; Photometrics). Optical z-sections were separated by 0.125 µm. Exposure times were between 3 and 10 ms, with three rotations of the illumination grid. Multichannel imaging was achieved through sequential acquisition of wavelengths by separate cameras. First, the channels were aligned in the image plane and around the optical axis using predetermined shifts and measured using a target lens and the SoftWoRx alignment tool. Afterwards, they were carefully aligned using alignment parameters from control measurements made with 0.5 µm diameter multi-spectral fluorescent beads. Raw 3D-SIM images were processed and reconstructed using the DeltaVision OMX SoftWoRx software package. The final voxel size was 40 nm x 40 nm x 125 nm.

### Gentamicin protection assay

Blood from infected animals (day 14 post infection) or over night *in-vitro* infected erythrocytes (see above) were incubated for 2 h at 35°C, 5% CO2 either in PBS alone or in PBS containing 40 µg/ml gentamicin. Cells were washed 3 times with PBS and centrifugation at 300 x g for 5 min. Cell pellets were frozen at −80°C for erythrocyte lysis and plated in serial dilutions on blood agar plates. Supernatant and medium only controls were directly used for serial dilution and plating.

### Antibody binding to the bacterial surface by flow cytometry

Bacteria were grown as described above, collected in PBS and stained with purified antibody for 45 min at RT. Bacteria were then washed twice with PBS, centrifuged at 4000 x g for 5 min and stained with the secondary antibody (anti-mouse IgG-Alexa 647) for 1 h at 4°C in the dark. After 3 washes, the bacteria were fixed in 3.7% PFA for 10 min at 4°C in the dark and resuspended in FACS buffer before analysis via flow cytometry (CantoII).

For CFA expression from plasmid, the bacteria were incubated in DMEM containing 10% FCS and 100 nM IPTG overnight prior to staining.

For binding curves, a serial dilution series of monoclonal antibody or isotype was performed in a U-bottom 96-well plate. 10^7^ bacteria were added per well. After 45 min at RT in the dark, staining continued as described above.

### ELISA

Purification of *Bartonella* outer membrane proteins (OMP) was performed as has been described previously (Otsuyama, et al., 2016). For antibody titers in serum, anti murine IgG-HRP or anti murine IgM-biotin and streptavidin-HRP were used.

In brief, ELISA plates were coated over night with purified *Bartonella* OMP (1 mg per 96-well plate) in coating buffer. The plates were washed with 0.05 % Tween in PBS and blocked 1 h at RT with 1x ELISA buffer. After another wash, sera were added in dilution series for 3 h at RT. After 3 washes, detection antibody was added in the recommended dilution for 1 h at RT. If necessary streptavidin-HRP was added after another 3 washes for 30 min at RT. After 5 washes 1x ELISA substrate was added at RT for 5-10 min and stopped with 1 M phosphoric acid. Plates were read at 450 and 570 nm using a plate reader.

### Hybridoma production and handling

For hybridoma production the DiSH Kit (Enzo Life Sciences) was used. BALB/c mice were infected as described above before *i.v.* reinfection with 10^7^ cfu after clearance of the first infection. 2 weeks later, the animals were boosted with 10^7^ cfu heat inactivated bacteria (3 h at 60°C) and the spleens were harvested 2 days later. A collagenase digest (3 mg/ml Collagenase IV, 2% FCS, in RPMI) was performed for 30-60 min at 37°C. The digestion mixture was put through a cell strainer and approx. 10 ml of RPMI + 10% FCS were added. The cells were centrifuged for 10 min at 300 x g at 4°C and a red blood cell lysis was performed by resuspending the pellet in ACK buffer (150 mM NH_4_Cl, 10 mM KHCO_3_ and 0.1 mM EDTA). After 1 min at RT, 10 ml of RPMI + 10% FCS were added and the cells were centrifuged again and used according to the DiSH Kit protocol. In brief, the obtained lymphocytes were washed with WCM. Myeloma fusion partner SP2ab grown in PMC medium, washed once with WCM and added to the lymphocyte pellet in a ration of 1:5 Sp2ab: lymphocyte. After centrifugation for 10 min at 4°C, 300 x g, the cell pellet was warmed to 37°C in a water bath. While keeping the pellet in the water bath, PEG1000 was added drop wise to the cell mixture. After 60 sec at 37°C, 4 ml of prewarmed WCM were added drop-wise over the course of 2 min while keeping the cells at 37°C. Another 5 ml were added over the course of 1 min. Another 6 ml were added over the course of 1 min. After 10 min at 37°C, 30 ml of FCM were added and the cells were centrifuged at 150 x g for 5 min. The cells were then incubated over night in FCM at 37°C, 5% CO2. The fused cells were plated in semisolid medium complemented with hybridoma selection medium until the appearance of single colonies, approx. after 8 days. Clones were then harvested in RCM, expanded in ECM and screened for specificity using the EAI assay. Hybridoma clones were then grown in RPMI complemented with 10% FCS.

### Purification of monoclonal antibodies

Sequencing of the antibody expressed by a hybridoma cell line was performed by Absolute Antibody, UK. The sequences of LS4G2 and LS5G11 V_L_ and V_H_ region are given in Table S4.

The corresponding gene fragments were synthesized (Genscript, New Jersey, USA). For recombinant expression, rearranged V regions were subcloned into the CMV-promoter-driven mammalian expression vector pXLG1.2, followed by the Cγ2a constant domain (kindly provided by Prof. Shozo Izui, University of Geneva), corresponding to the Genbank sequences for mouse IgG2a (J00470.1). The procedure for the light chain was identical.

Antibodies were expressed by transient co-transfection of HEK cells (Protein Production and Structure Core Facility, EPFL, Lausanne, Switzerland) and purified on protein G columns with ÄKTAprime plus followed by PBS dialysis.

For conjugation of the purified antibody, we used the lightning link labeling kit from Expedeon according to the manufacturer’s protocol.

### Immunoprecipitation

*Bartonella taylorii* IBS296 was grown as described above. Bacteria were collected in wash buffer (50 mM HEPES pH 7.4, 200 mM NaCl, 5% glycerol) containing benzoase and protease inhibitors, lyzed using French Press and centrifuged at 10.000 x g for 10 min at 4°C. The supernatant was centrifuged at 100.000 x g at 4°C to obtain the membrane protein fraction. The pellet was solubilized over-night at 4°C with 1% DDM in wash buffer and again centrifuged at 100.000 x g at 4°C. The supernatant was incubated overnight with 20 µg of antibody and 80 µl of UltraLink Resin at 4°C. The mixture was centrifuged at 6000 x g and the beads were washed 3 times with excess wash buffer containing 0.05% DDM before elution with 25 µl of Laemmli buffer. If indicated the samples were digested using 1 U or proteinase K for 10 min at 4°C before another 3 washing steps and elution.

Sample preparation for MS-based proteome analysis 0.5 µL of 1 M iodoacetamide was added to the samples. Cysteine residues were alkylated for 30 min at 25°C in the dark. Digestion and peptide purification was performed using S-trapTM technology (Protifi) according to the manufacturer’s instructions. In brief, samples were acidified by addition of 2.5 µL of 12% phosphoric acid (1:10) and then 165 µL of S-trap buffer (90% methanol, 100 mM TEAB pH 7.1) was added to the samples (6:1). Samples were briefly vortexed and loaded onto S-trapTM micro spin-columns and centrifuged for 1 min at 4000 g. Flow-through was discarded and spin-columns were then washed 3 times with 150 µL of S-trap buffer (each time samples were centrifuged for 1 min at 4000 g and flow-through was removed). S-trap columns were then moved to the clean tubes and 20 µL of digestion buffer (50 mM TEAB pH 8.0) and trypsin (at 1:25 enzyme to protein ratio) were added to the samples. Digestion was allowed to proceed for 1 h at 47°C. After, 40 µL of digestion buffer was added to the samples and the peptides were collected by centrifugation at 4’000 g for 1 minute. To increase the recovery, S-trap columns were washed with 40 µL of 0.2% formic acid in water (400g, 1 min) and 35 µL of 0.2% formic acid in 50% acetonitrile. Eluted peptides were dried under vacuum and stored at −20 °C until further analysis.

The peptides were dissolved in LC-buffer A (0.15% formic acid, 2% acetonitrile) right before ultrasonication for 10 sec and shaking at 1’400 rpm at 25°C for 5 min.

### Mass spectrometry

For each sample, aliquots of 0.4 μg of total peptides were subjected to LC-MS analysis using a dual pressure LTQ-Orbitrap Elite mass spectrometer connected to an electrospray ion source (both Thermo Fisher Scientific) and a custom-made column heater set to 60°C. Peptide separation was carried out using an EASY nLC-1000 system (Thermo Fisher Scientific) equipped with a RP-HPLC column (75 μm × 30 cm) packed in-house with C18 resin using a linear gradient from 95% solvent A (0.1% formic acid in water) and 5% solvent B (80% acetonitrile, 0.1% formic acid, in water) to 35% solvent B over 50 minutes to 50% solvent B over 10 minutes to 95% solvent B over 2 minutes and 95% solvent B over 18 minutes at a flow rate of 0.2 μl/min. The data acquisition mode was set to obtain one high resolution MS scan in the FT part of the mass spectrometer at a resolution of 120’000 full width at half maximum (at 400 m/z, MS1) followed by MS/MS (MS2) scans in the linear ion trap of the 20 most intense MS signals. The charged state screening modus was enabled to exclude unassigned and singly charged ions and the dynamic exclusion duration was set to 30 s. The collision energy was set to 35%, and one microscan was acquired for each spectrum. The mass spectrometry proteomics data have been deposited to the ProteomeXchange Consortium via the PRIDE (Perez-Riverol, et al., 2019) partner repository with the dataset identifier PXD028783 and 10.6019/PXD028783.

### Protein Identification and Label-free Quantification

The acquired raw-files were imported into the Progenesis QI software (v2.0, Nonlinear Dynamics Limited), which was used to extract peptide precursor ion intensities across all samples applying the default parameters. The generated mgf files were searched using MASCOT against a decoy database containing normal and reverse sequences of *B. taylorii* IBS296 (UniProt, 10.01.2020) proteome and commonly observed contaminants (in total 3484 sequences) generated using the SequenceReverser tool from the MaxQuant software (Version 1.0.13.13). The following search criteria were used: full tryptic specificity was required (cleavage after lysine or arginine residues, unless followed by proline); 2 missed cleavages were allowed; carbamidomethylation (C) was set as fixed modification; oxidation (M) and protein N-terminal acetylation were applied as variable modifications; mass tolerance of 10 ppm (precursor) and 0.6 Da (fragments) was set. The database search results were filtered using the ion score to set the false discovery rate (FDR) to 1% on the peptide and protein level, respectively, based on the number of reverse protein sequence hits in the datasets. Quantitative analysis results from label-free quantification were normalized and statically analyzed using the SafeQuant R package v.2.3.4 to obtain protein relative abundances. This analysis included summation of peak areas per protein and LC MS/MS run followed by calculation of protein abundance ratios. Only isoform specific peptide ion signals were considered for quantification. The summarized protein expression values were used for statistical testing of differentially abundant proteins between conditions. Here, empirical Bayes moderated t-Tests were applied, as implemented in the R/Bioconductor limma package. The resulting p-values were adjusted for multiple testing using the Benjamini Hochberg method.

### DNA extraction and sequencing

Genomic bacterial DNA was extracted with QIAGEN Genomic-tip 20/G according to the manufacturer’s guidelines. Whole genome sequencing of *B. taylorii* M1 strain was done with Illumina and Nanopore sequencing technologies. Illumina short-read sequencing was performed at the Microbial Genome Sequencing Center (MiGS) using the Illumina NextSeq 2000 platform. The libraries for the Nanopore sequencing were made with Ligation Sequencing Kit and Native Barcoding Expansion 1-12 according to the protocol provided by ONT. The sequencing was done with the MinION Mk1B sequencing device and the MinION Flow Cell (R10.3).

### Genome assembly and annotation

All calculations were performed at sciCORE scientific computing center at the University of Basel. Base- calling and barcoding of raw ONT data was done with Guppy v.5.0.7 (Oxford Nanopore Technologies). Then, ONT read sets were filtered and reads shorter than 1 kbp were removed with Filtlong v.0.2.0. Illumina reads were quality assessed with FastQC v.0.11.8 and if needed trimmed with Trimmomatic v.0.39. Next, consensus assemblies were generated with Trycycler v.0.5.0 based on multiple input assemblies made with Canu v.2.1.1, Raven v.1.5.0, Minimap2 v.2.20-r1061 combined with Miniasm v.0.3-r179 and polished with Minipolish v.0.1.3. The assemblies then were finalised by polishing with ONT reads (Medaka v.1.4.3) and short reads (Pilon v.1.24).

### Bioinformatic analysis

Synteny analysis of Cfa autotransporter was inferred with MultiGeneBlast v.1.1.14 in architecture search mode with default search parameters. The database was generated in MultiGeneBlast from annotated genome sequences of *Bartonella* species (Table S1).

Search for domains present in CFA was done with Conserved Domain Search tool with Result mode set on “Full”.

The alignment of protein sequences of Cfa was made with MSAProbs v.0.9.7 implemented in MPI Bioinformatics Toolkit. Positions of the alignment containing at least 40% of gaps were removed from the alignment with Mask alignment option of Geneious Prime. Resulting trimmed alignment was used to calculate conservation scores with the Property Entropy (PE) scoring method of Protein Residue Conservation Prediction tool. Conservation scores were summed and averaged within a sliding window of 19 amino acids [ai-9; ai+9], where a is a conservation score and i is an amino acid index. The resulting values were plotted with GraphPad Prism. Alignment of Cfa from lineage 4 species was used to identify variability sites. Sites were considered variable if averaged PE scores were equal or lower 0.4. Then, variability sites were mapped onto Cfa from *B. taylorii* IBS296 and alignments generated for *B. quintana*. Modelling the Cfa from *B. taylorii* IBS296 was done with AlphaFold with default options.

Tree topology and bootstrap support values were inferred with PhyML v.3.0, model (JTT+G+I+F), 1000 bootstrap replicates. The tree was based on a concatenated alignment of five core protein sequences (FtsZ, GroEL, GyrB, RpoB and SecY). The protein sequences were aligned with Clustal Omega (implemented in Geneious Prime), then concatenated, and all gaps were removed with Mask alignment option of Geneious Prime.

### Data analysis

Unless stated differently, statistical analysis of the obtained data was performed using GraphPad Prism Software and the statistical tests as indicated. P-values are depicted as follows: ns = P > 0.05; *= P ≤ 0.05; ** = P ≤ 0.01; *** = P ≤ 0.001 All LC-MS analysis runs were acquired from independent biological samples. To meet additional assumptions (normality and homoscedasticity) underlying the use of linear regression models and Student t-Test MS-intensity signals are transformed from the linear to the log-scale. Unless stated otherwise linear regression was performed using the ordinary least square (OLS) method as implemented in base package of R v.3.1.2. The sample size of three biological replicates was chosen assuming a within-group MS-signal Coefficient of Variation of 10%. When applying a two-sample, two-sided Student t test this gives adequate power (80%) to detect protein abundance fold changes higher than 1.65, per statistical test. Note that the statistical package used to assess protein abundance changes, SafeQuant, employs a moderated t-Test, which has been shown to provide higher power than the Student t-test. We did not do any simulations to assess power, upon correction for multiple testing (Benjamini-Hochberg correction), as a function of different effect sizes and assumed proportions of differentially abundant proteins. For peptide enrichment upon proteinase K digest, the ratio between digested and undigested sample was formed and plotted within a sliding window of 19 amino acids [ai-9; ai+9], where a is a conservation score and i is an amino acid index.

