## supplemental information for "The *Bartonella* autotransporter CFA is a protective antigen and hypervariable target of neutralizing antibodies blocking erythrocyte infection"

**Figure S1 (Related to Figures 1 and 2)**

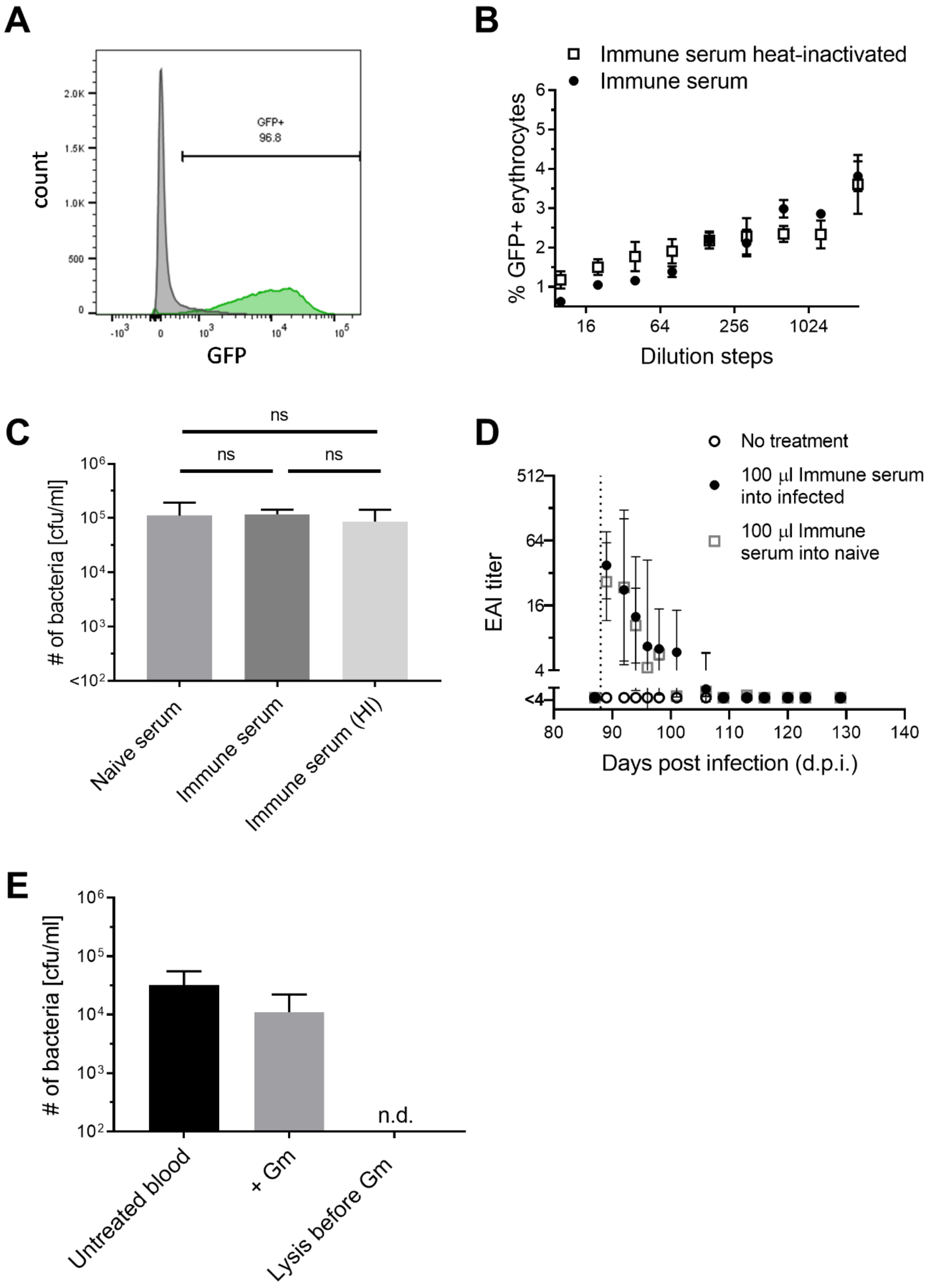


**Figure S1**: **Complement-independent EAI activity and lack of bactericidal effect of antisera and gentamicin-insensitivity of intraerythrocytic Bartonella. (A)** Flow cytometry analysis of wild-type (WT) and GFP-expressing B. taylorii IBS296. **(B)** EAI assay comparing untreated immune serum and heat-inactivated (60°C, 30 min) immune serum. **(C)** Bacterial survival in the EAI assay. Bacteria were plated at the end of an EAI assay comparing the adhesion-inhibiting activity of naïve serum, anti-B. taylorii IBS296-immune serum and heat-inactivated anti-B. taylorii IBS296-immune serum. **(D)** Decay of EAI titer in the serum of B-cell knock-out mice receiving immune serum transfer in the experiment reported in Figure 1C, D (100 µl immune serum into infected), side-by-side with titers determined in a separate group of uninfected B-cell knock-out mice that received the same serum transfer (100 µl immune serum into naïve) and a group of uninfected B-cell knock-out mice without serum transfer (no treatment). The vertical dotted line indicates the time point of serum transfer. A baseline sample was collected prior to serum transfer on d87. **(E)** Lysis control for the gentamicin (Gm) protection assay shown in Figure 2E. Blood was collected from infected wild-type (WT) mice on day 14 p.i.. Cfu/ml counts are shown for untreated blood samples, Gm-treated erythrocytes from infected animals and erythrocytes which had been lysed by a freeze-thaw cycle prior to Gm treatment. All data show representative results from two independent experiments; in vitro experiments were performed in technical triplicates and in vivo experiments with at least 3 mice per group and experiment. Data are displayed as mean ± SD. Statistical analysis was performed using one-way ANOVA, ns: $P > 0.05$; HI = heat inactivated.

**Figure S2 (Related to Figure 3)**

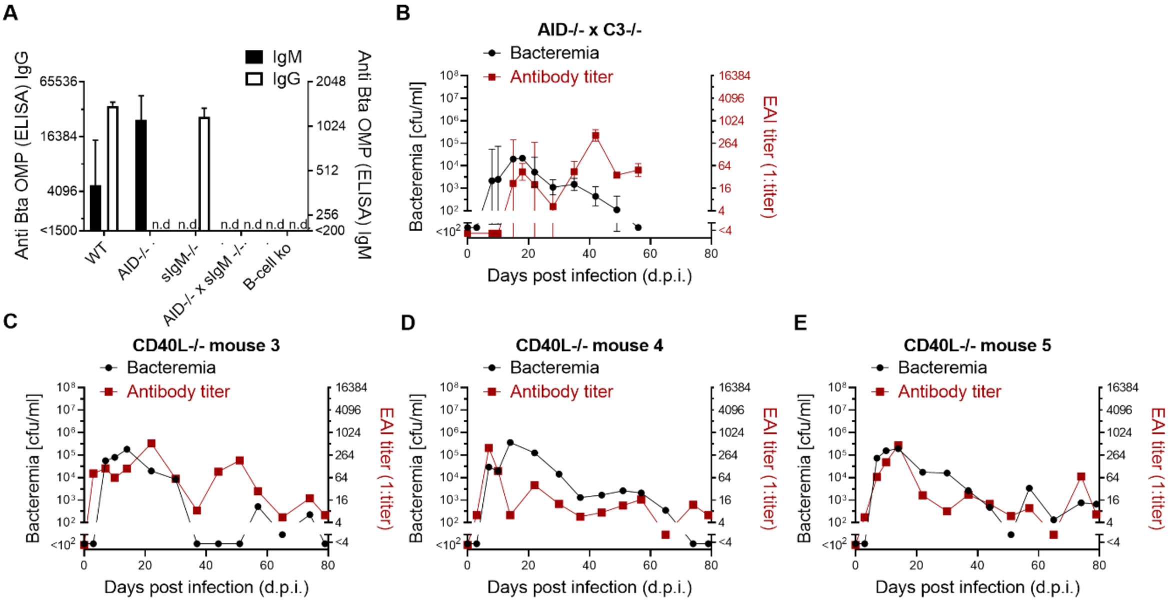


**Figure S2**: **Antibody response and *B. taylorii* IBS296 bacteremia in mice lacking AID, soluble IgM, C3 and CD40L or combinations thereof. (A)** IgM (right Y-axis) and IgG (left Y-axis) antibodies binding to the *B. taylorii* IBS296 (Bta) outer membrane proteins (OMPs) fraction were measured by ELISA. Sera were collected from wild-type (WT), AID-/-, sIgM-/- -, AID-/- x sIgM-/- and B-cell knock-out mice on day 42 days after infection. **(B)** We infected AID-/- x C3 -/- mice with *B. taylorii* IBS296 and measured bacteremia and serum EAI titer over time. **(C-E)** Bacteremia and EAI serum titers of individual CD40L-/- mice. Each plots represents one single animal from the experimental group reported in Figure 3G. **(A-B)** Representative data from two independent experiments with at least four mice per group are shown. **(A, B)** Data are represented as mean ± SD; n.d. = not detected.

### Figure S3 (Related to Figures 4 and 5)

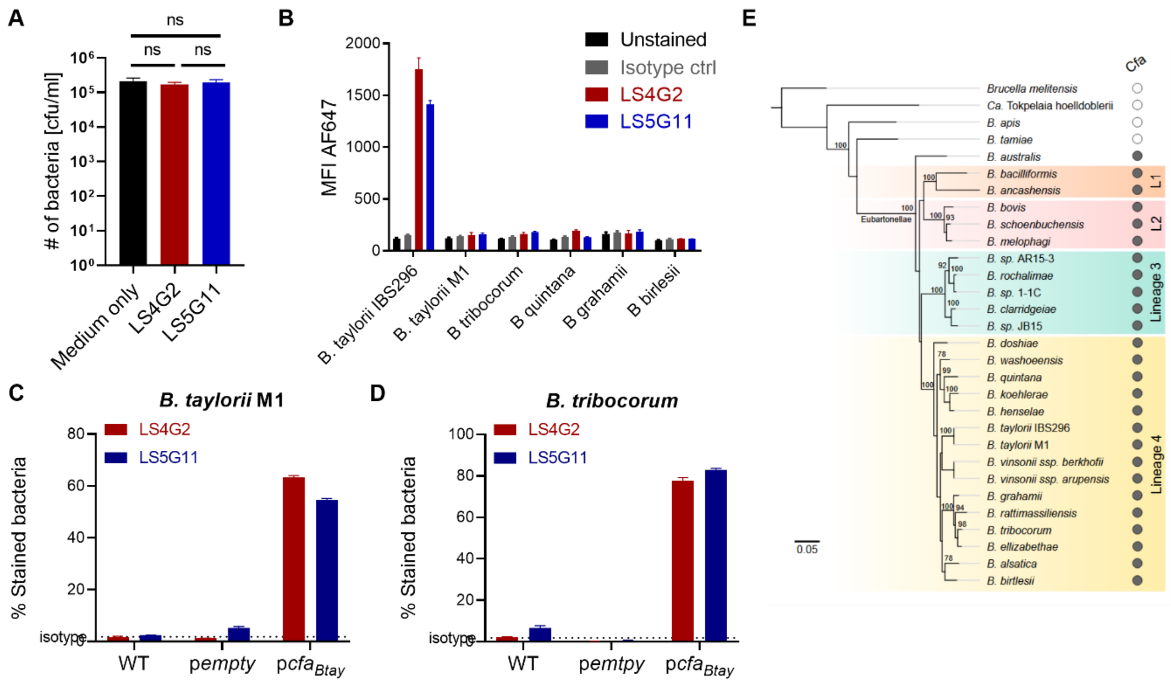


**Figure S3: Characterization of monoclonal antibodies targeting CFA and conservation of this autotransporter within the Bartonellae. (A)** Survival of *B. taylorii* IBS296 in the EAI assay. Bacteria were incubated with either one of the monoclonal antibodies LS4G2 and LS5G11 or with medium control. Surviving bacteria were quantitated 24 h later. **(B)** Staining of *B. taylorii* IBS296, *B. quintana*, *B. tribocorum*, *B. taylorii* M1, *B. grahamii* and *B. birtlesii* with LS4G2 (red), LS5G11 (blue) or irrelevant isotype control antibody (grey) was compared by flow cytometry. Unstained bacteria of each species (black) served as background control. **(C, D)** Flow cytometry staining of isogenic strains of **(C)** *B. taylorii* M1 or **(D)** *B. tribocorum*, either in their unmodified wild-type (WT) form or transformed with either empty vector control (p*empty*) or plasmid p*cfa*$_{Btay}$ driving expression of CFA from *B. taylorii* IBS296. Bars show the percentage of bacteria stained with LS4G2 or LS5G11, respectively, as determined by flow cytometry. All data show representative results from three independent experiments, which were performed in technical triplicates. Data are represented as mean ± SD. Data presented in (A) were analysed 1-way ANOVA. ns = P > 0.05. **(E)** Phylogeny of the *Bartonella* genus and the related *Brucella melitensis* and "*Candidatus Tokpelaia hoelldoblerii*" as outgroup taxons. Filled and empty circles show the presence or absence, respectively, of genes encoding the CFA autotransporter. Colouring scheme represents lineages of *Bartonella*, orange (L1), red (L2), green (L3) and yellow (L4). Values above nodes show bootstrap support (>75%). The scale bar indicates the number of substitutions per site.

### Figure S4 (Related to Figure 6)

### A

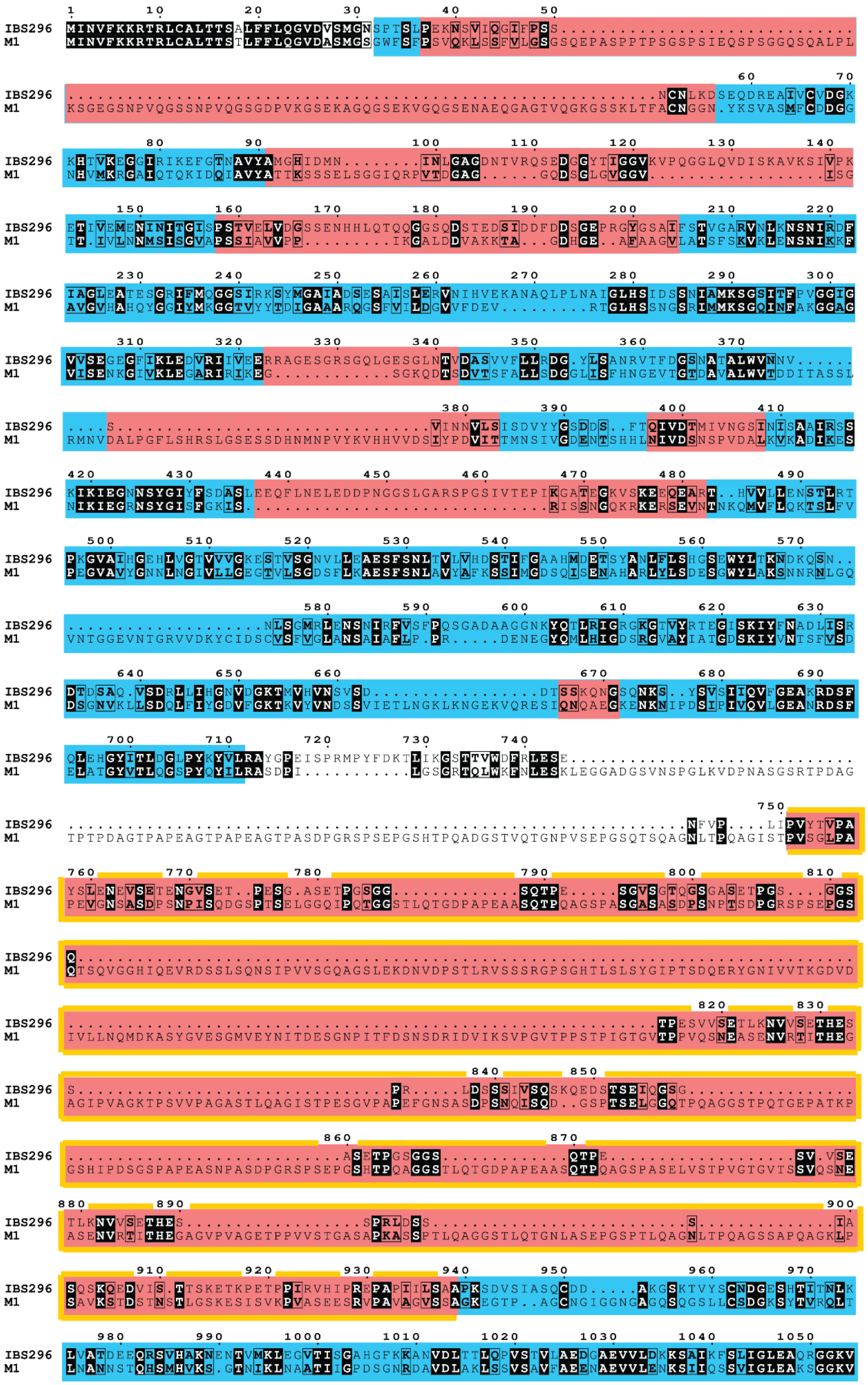

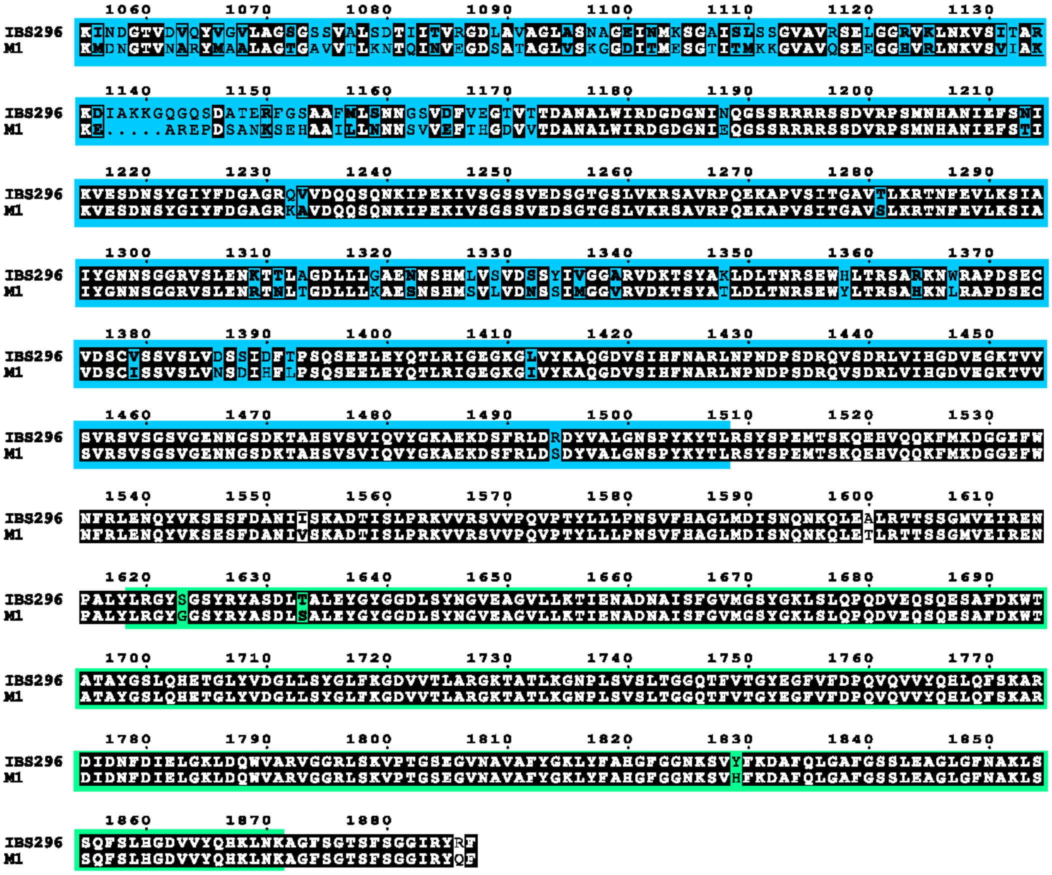

B

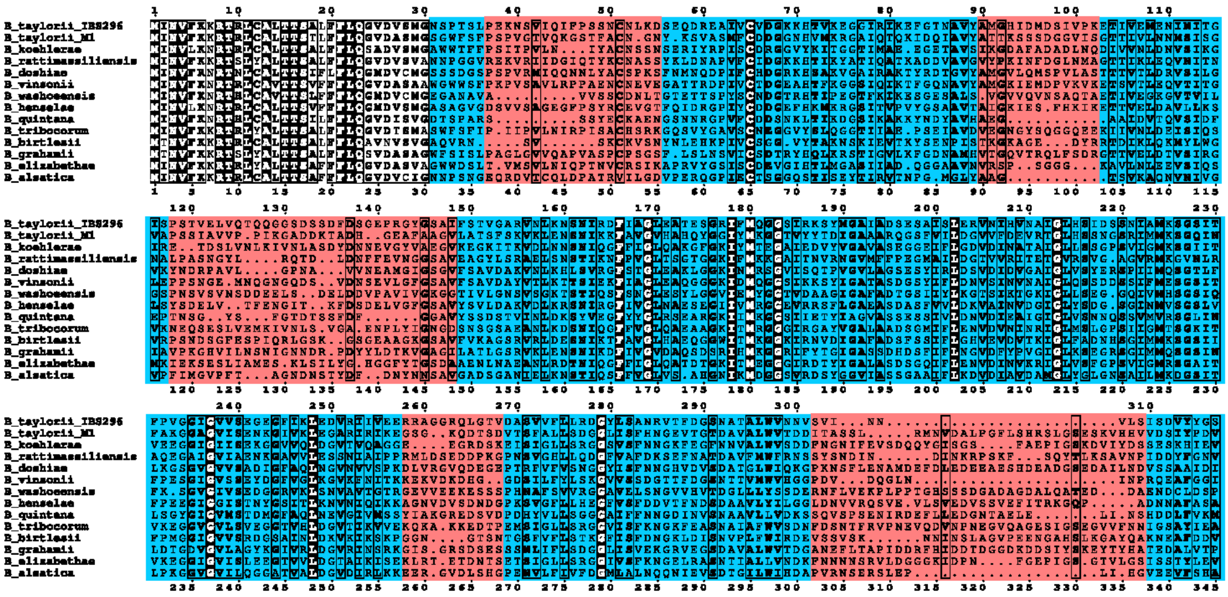

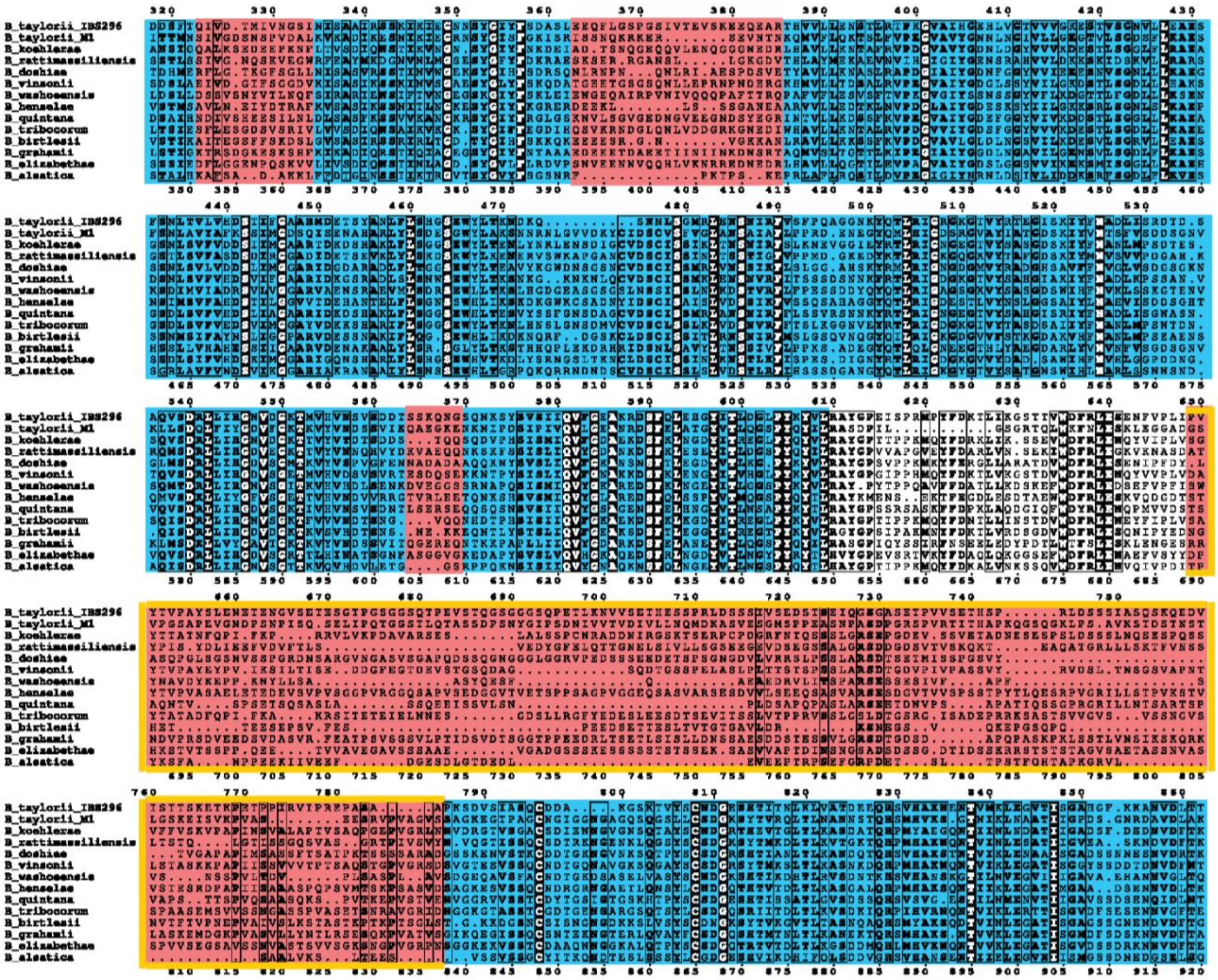

80

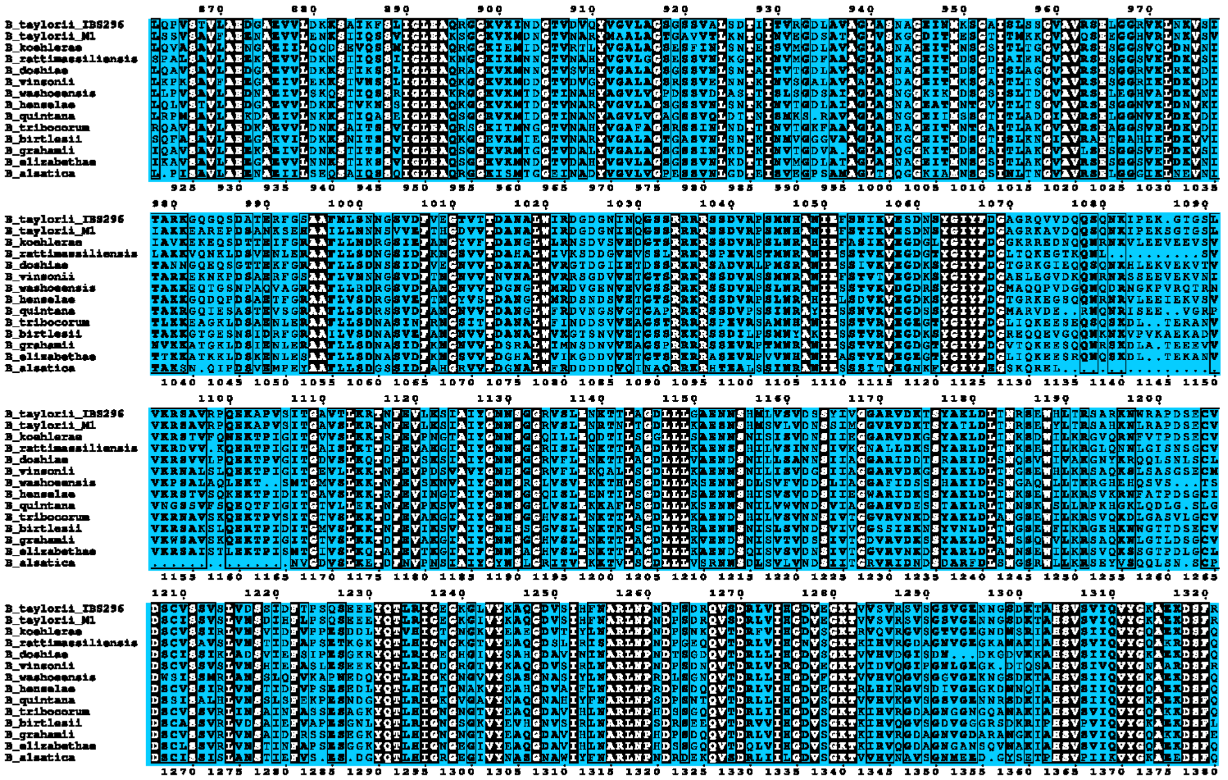

81

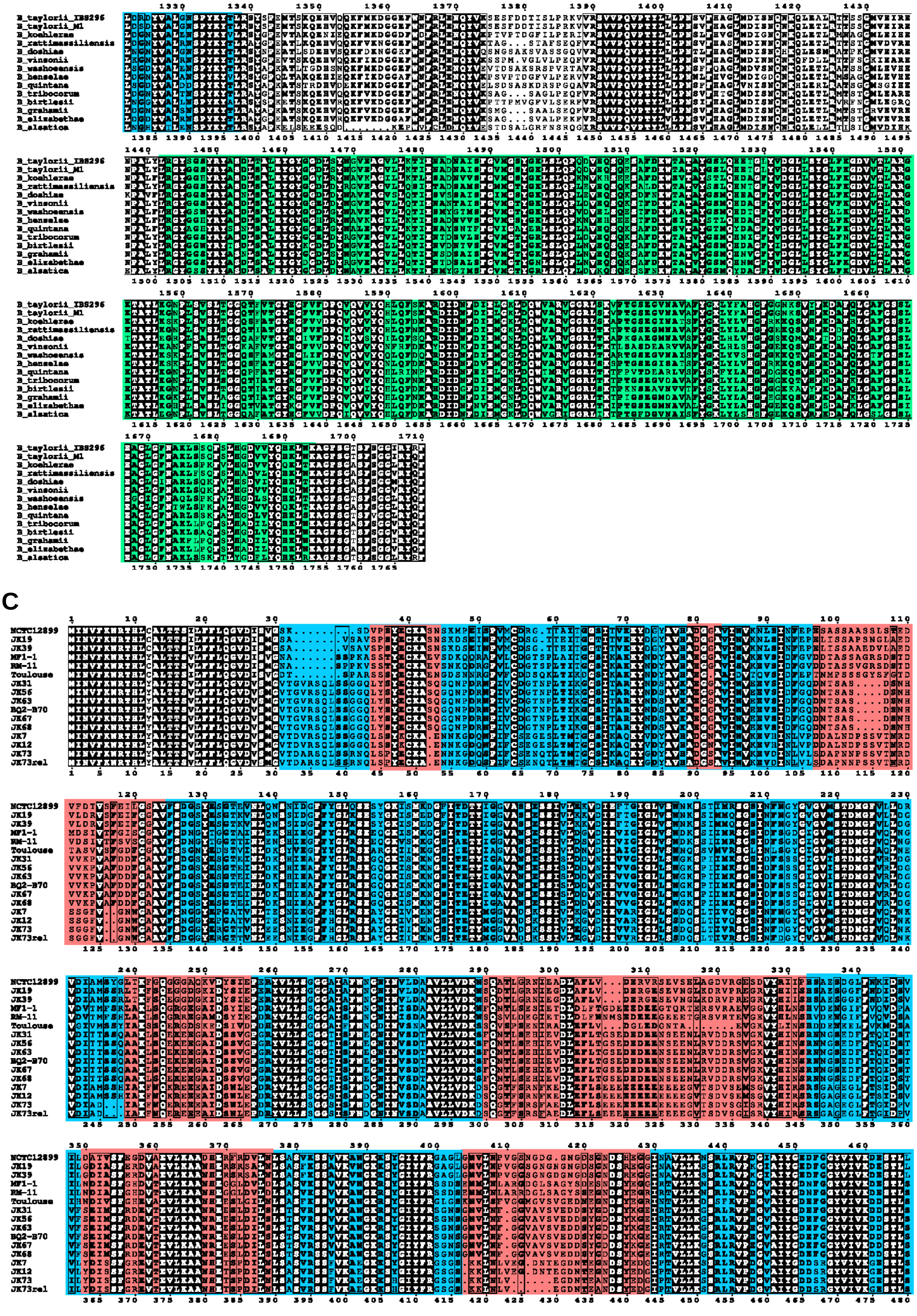
B_taylorii_IBS296
B_taylorii_M1
B_koehlerae
B_rattimassiliensis
B_doshiae
B_vinsonii
B_washoensis
B_henselae
B_quintana
B_tribocorum
B_birtlesii
B_grahamii
B_elizabethae
B_alsatica
C
NCTC12899
JK19
JK39
MF1-1
RM-11
Toulouse
JK31
JK56
JK63
BQ2-B70
JK67
JK68
JK7
JK12
JK73
JK73rel

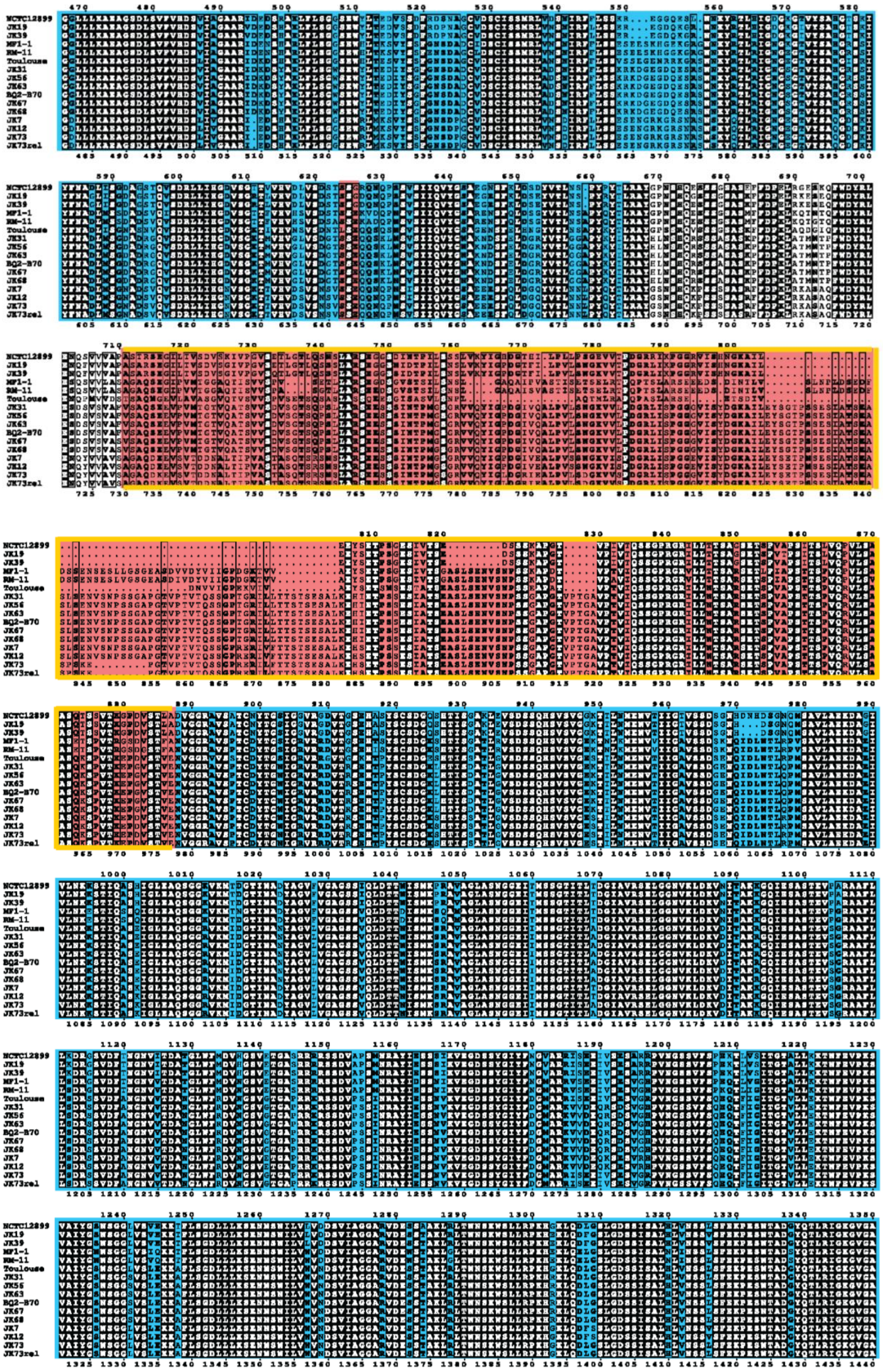
NCTC12899
JK19
JK39
MF1-1
RM-11
Toulouse
JK31
JK56
JK63
BQ2-B70
JK67
JK68
JK7
JK12
JK73
JK73rel

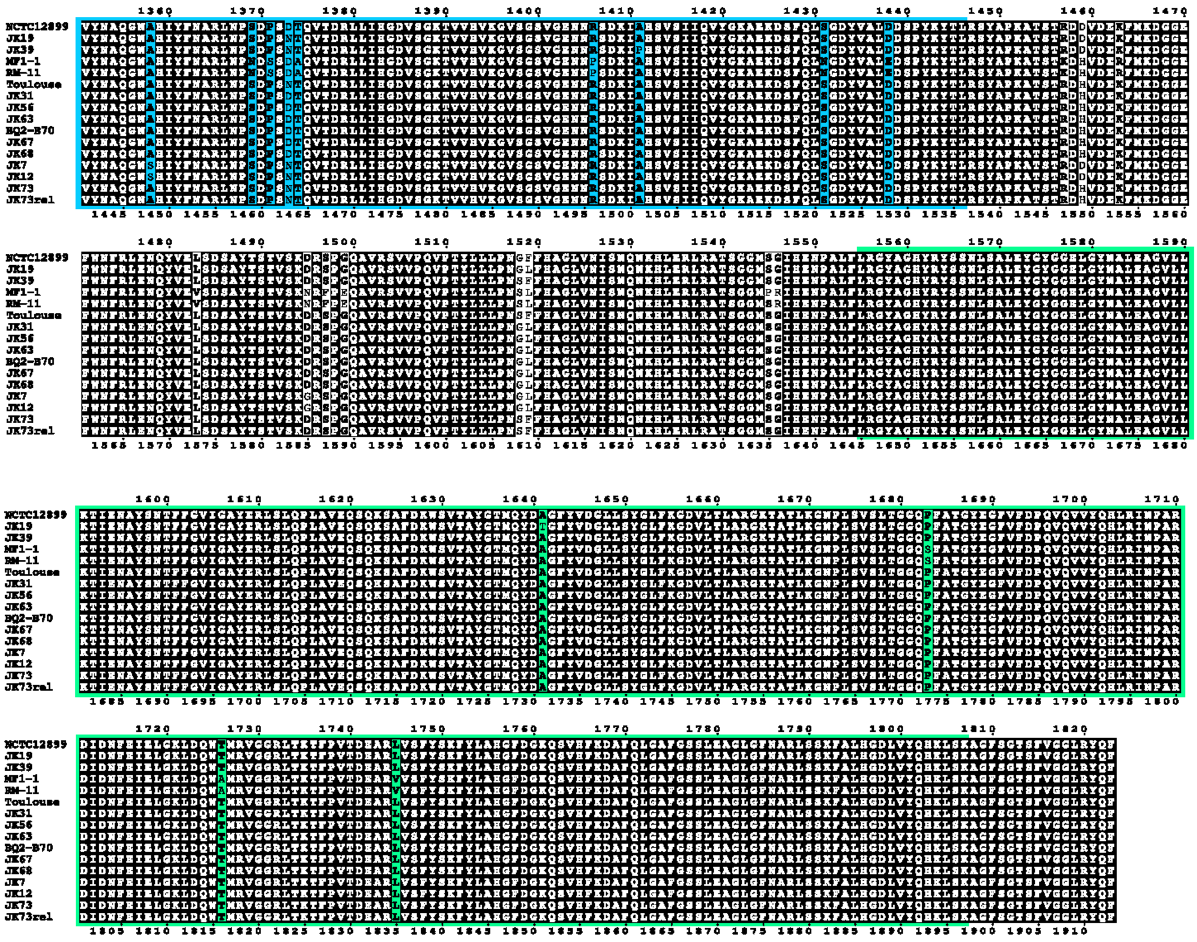

**Figure S4**: **Sequence variation in CFA between and within individual *Bartonella* lineages and species.** Protein sequence alignments for **(A)** *B. taylorii* IBS296 and M1, another *B. taylorii* isolate, **(B)** representative species of *Bartonella* lineage 4 and **(C)** a collection of clinical isolates of *B. quintana*. Black indicates high sequence conservation, β-helices (stalk) are labelled in blue and in the β-barrel in green. Hyper-variable regions are highlighted in red and the linker between β-helices is additionally highlighted in orange.

**Figure S5 (Related to Figure 6)**

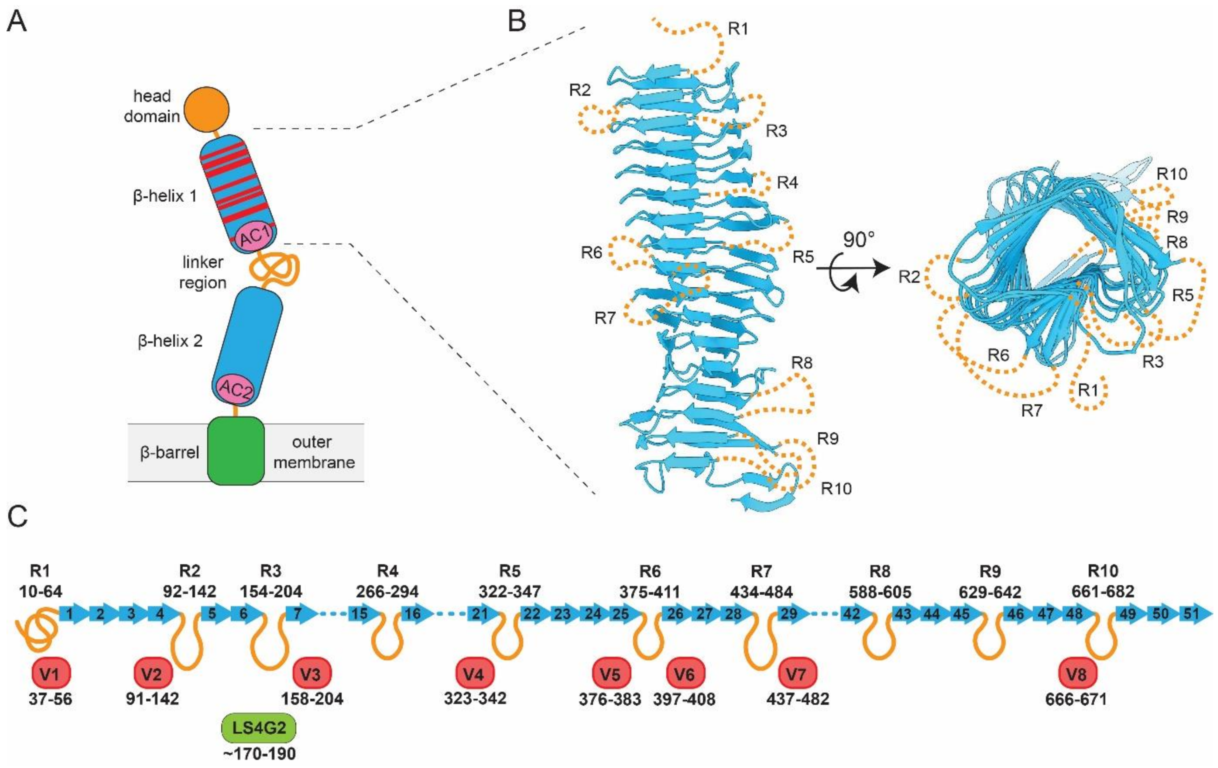


**Figure S5**: **Structure prediction for the N-terminal stalk domain of CFA. (A)** Schematic overview of the CFA domain topology. The N-terminal head domain (orange) is followed by two β-helices (blue) connected by an unstructured linker. A C-terminal β-barrel (green) anchors the structure to the outer membrane. Red lines indicate variability regions in the N-terminal β-helix as determined by sequence alignment. AC, autochaperone domains. **(B)** A three-dimensional structure prediction of CFA's β-helix 1 from *B. taylorii* IBS296 was generated using AlphaFold2. The model indicates a conserved pertactin-like fold with a rigid core formed by several turns of β-strands (blue). The modeling approach indicated a number of regions that likely fall out of register of the β-helix, which suggests that they protrude to the outside and decorate the stalk (R1-R10, dotted orange lines). **(C)** Schematic representation of the AlphaFold2 model shown in (B). 51 β-strands forming the core of the helix are shown as blue arrows. R1-R10 regions are indicated as orange loops. The locations of the hypervariability regions (V1-V8) are shown in red. The location of the putative LS4G2 binding site is depicted in green. Amino acid residues comprised in the respective regions are indicated.

**Table S1:** Accession numbers of the sequences used in this study.

| Species | Strain | NCBI Accession Number |
|---|---|---|
| *Bartonella alsatica* | CIP 105477 | NZ_CP058235 |
| *B. ancashensis* | 20.00 | NZ_CP010401 |
| *B. apis* | PEB0149 | NZ_LXYT01000001 |
| *B. australis* | Aust/NH1 | NC_020300 |
| *B. bacilliformis* | KC583 | NC_008783 |
| *B. birtlesii* | LL-WM9 | NZ_AIMC00000000.1 |
| *B. bovis* | 91-4 | NZ_CM001844 |
| *B. clarridgeiae* | 73 | NC_014932 |
| *B. doshiae* | NCTC12862 | NZ_UFTF01000001 |
| *B. elizabethae* | NCTC12898 | NZ_LR134527 |
| *B. grahamii* | as4aup | CP001562 |
| *B. melophagi* | K-2C | NZ_AIMA00000000.1 |
| *B. henselae* | Houston-1 | NC_005956 |
| *B. koehlerae* | C-29 | NZ_AHPL00000000.1 |
| *B. quintana* | Toulouse | NC_005955 |
| | NCTC12899 | NZ_LS483373 |
| | JK19 | NZ_AHPH00000000.1 |
| | JK39 | NZ_AHPB00000000.1 |
| | MF1-1 | NZ_AP019773 |
| | RM-11 | NC_018533 |
| | JK31 | NZ_AHPG00000000.1 |
| | JK56 | NZ_AHPE00000000.1 |
| | JK63 | NZ_AHPF00000000.1 |
| | BQ2-B70 | NZ_AZZV00000000.1 |
| | JK67 | NZ_AHPC00000000.1 |
| | JK68 | NZ_AHPD00000000.1 |
| | JK7 | NZ_AZZZ00000000.1 |
| | JK12 | NZ_AZZY00000000.1 |
| | JK73 | NZ_AZZX00000000.1 |
| | JK73rel | NZ_AZZW00000000.1 |
| *B. rattimassiliensis* | 15908 | NZ_AILY00000000.1 |
| *B. rochalimae* | BMGH | NZ_AHPK00000000.1 |
| *B. schoenbuchensis* | R1 | NZ_CP019789 |
| *B. sp.* | 1-1C | NZ_CP019489 |
| | AR15-3 | NZ_MUYE00000000 |
| | JB15 | NZ_CP019787 |
| *B. taylorii* | IBS296 | |
| | M1 | |
| *B. tamiae* | Th239 | NZ_AIMB00000000 |
| *B. tribocorum* | CIP 105476 | NC_010161 |
| *B. vinsonii* | NCTC12905 | NZ_LR134529 |
| *B. vinsonii ssp. arupensis* | Pm136co | NZ_AIMH00000000.1 |
| | OK-94-513 | NZ_AILZ00000000.1 |
| *B. vinsonii ssp. berkhofii* | Winnie | NC_020301.1 |
| | Tweed | NZ_AGWD00000000.1 |
| *B. washoensis* | Sb944nv | NZ_AILU00000000.1 |
| "*Candidatus* Tokpelaia hoelldoblerii" | Hsal | CP017315 |
| *Brucella melitensis* | 16M | AE008917, AE008918 |

**Table S2: List of oligonucleotides used in this study**

| fw primer | rev primer | amplified target | purpose |
|---|---|---|---|
| prJS370: 5'CTCATCCTGTCTCTTGATCAGATC | prJS371: 5'AGATCTGGGGTTCGAAATGACCG | pJC43 excluding KanR | Generation of pLS04 |
| prJS372: 5'TCAAGAGACAGGATGAGAAGCCCTGCAAAGTAAACTGG | prJS373: 5'TTTCGAACCCCAGATCTTTAGGTGGCGGTACTTGGG | Gentamycin cassette of pPG1000 | Generation or pLS04 |
| prLS213: 5'CATCCTGACGCCCCCGGGGATCCTAGCTTCGTTAAGTCCAGTGAGACC | prLS214: 5'GTCAAACTTTCTAAATATAATTTGCATAAAGAA | Homology region 1 *cfa* locus of LSB001 | Generation or pLS25 |
| prLS230: 5'ATGCAAATTATATTTAGAAAGTTTGACATCTGCGATGATAGCTATTGTTGG | prLS231: 5'TGCAGAGCTTAGCTCTGCAGGTCGACGAATAAACCAACCTTCGTCTCCG | Homology region 2 *cfa* locus of LSB001 | Generation or pLS25 |
| prLS213 | prLS231 | SOEing PCR homology region 1 and 2 *cfa* locus | Generation or pLS25 |
| prJS1003: 5'GGCCTATTAAGACCCACTCACTGCCCGCTTTCCAG | prJS1004: 5'CATTGATCATATTGTTCCTCCTATGAGCTCTAGAGG | Amplification of regulatory region including lacI and P*lac* of pBZ485 | Generation of pLS51 |
| prJS1005: 5'GAGGAACAATATGATCAATGTATTTAAAAAGCGTACACG | prJS1006: 5'CTTCCATGTCGGCAGAACTAGAAACGATAGCGGATTCCG | Amplification of *cfa* or LSB001 | Generation of pLS51 |
| prJS1007: 5'TTCTGCCGACATGGAAGC | prJS893: 5'GTGGGTCTTAATAGGCCGACTGCGATG | Amplification of JC43 backbone without insert and promotor | Generation of pLS51 |
| prLS363: 5'CTTCCATGTCGGCAGAAGTCGACATTGTTCCTCCTATGAGCTCTAG | prLS364: 5'ATAGGAGGAACAATGTCGACTTCTGCCGACATGGAAGC | pLS51 without *cfa* insert | Generation of pLS60 |
| prJS1187: 5'ACCTTAAATGGATATATCG | prJS1188: 5'ACCAATAACTGCCT | Amplification of *cfa* including ist | Generation or pLS80 |

| | | | |
|---|---|---|---|
| CAACAACCAAAACAGAAGTTACCC | TAAACGTGATTACACTAGAAACGATAGCG | putative natural promotor from LSB001 | |
| prJS436: 5’ GGAGCTTGCGGCCCGGACGAATTCTCAGAAGAACTCGTCAAGAAGG | prJS478: 5’GGTTTATTAAATGATTGAACAAGATGGATTGC | Amplification of KanR of JC43 | Generation or pJS208 |
| prJS430: 5’ CCATTTAAGGTGATAGGTAAGATTATACC | prJS749: 5’GTTCAATCATTTAATAAACCTCCTTTCGGATCCG | Amplification of P*aphT* of JC43 | Generation of pJS208 |
| prJS428: 5’CACCAATAACTGCCTTAAAAGCTTATATATATGAGCTCTTCACTTTTCTCTATCACTGATAGG | prJS429: 5’ACCTATCACCTTAAATGGATATATCCCGGGATATATTTAAGACCCACTTTCACATTTAAG | Amplification of *tetR*-P*tetR* from synthetic gene fragment | Generation of pJS208 |

**Table S3: List of plasmids used in this study**

| plasmid | purpose | resistance | Cloning strategy | generation | source |
|---|---|---|---|---|---|
| pBZ485 | Generation of pLS51 | kanamycin | . | - | (Harms, et al., 2017) |
| pXLG1.2 | Generation of pLS06, 09, 34 and 35 | ampicillin | - | - | (Backliwal, et al., 2008), kindly provided by Prof. Shozo Izui, University of Geneva |
| pJC43 | Generation of pLS04 | kanamycin | - | - | (Celli, et al., 2005) |
| pPG1000 | Generation of pLS04 | gentamycin | - | - | (Schülein, et al., 2005) |
| pTR1000 | Generation of pLS25 | kanamycin | - | - | (Schulein & Dehio, 2002) |
| pUC18T-mini-Tn7 | Generation of pJS208 | gentamycin | - | - | (Choi, et al., 2005) |

| | | | | | |
|---|---|---|---|---|---|
| pLS04 | Constant GFP expression in *B. taylorii* | gentamycin | In-fusion ligation | Exchange of the kanamycin resistance cassette in pJC43 with a gentamycin cassette (pPG1000) | This study |
| pLS06 | LS4G2 IgG2a heavy chain expression | ampicillin | Restriction digest using NotI and MfeI | Exchange of previous V-region of pXLG1.2 with synthesized gene fragment LS4G2 $V_H$ | This study |
| pLS09 | LS4G2 kappa light chain expression | ampicillin | Restriction digest using NotI and HindIII | Exchange previous V region of pXLG1.2 with synthesized gene fragment LS4G2 $V_L$ | This study |
| pLS25 | Generation of *B. taylorii Δcfa* | kanamycin | Restriction digest using SalI and BamHI | Insertion of homology regions 1 and 2 into pTR000 | This study |
| pLS34 | Expression of LS5G11 IgG2a heavy chain | ampicillin | Restriction digest using NotI and MfeI | Exchange of previous V-region of pXLG1.2 with synthesized gene fragment LS5G11 $V_H$ | This study |
| pLS35 | LS5G11 kappa light chain expression | ampicillin | Restriction digest using NotI and HindIII | Exchange previous V region of pXLG1.2 with synthesized gene fragment LS5G11 $V_L$ | This study |
| pLS51 | IPTG inducible expression of *cfa* in *Bartonella* | kanamycin | In-fusion ligation | Ligation of JC43 backbone with P*lac* of pBZ485 and *cfa* gene | This study |
| pLS60 | Empty vector control for pLS51 | kanamycin | In-fusion ligation | Amplification of pLS51 lacking *cfa* insert | This study |
| pLS80 | Insertion of *cfa* containing its natural promoter into *Bartonella* using Tn7 transposon | kanamycin, ampicillin | In-fusion ligation | *cfa* including its putative natural promotor was ligated into XmaI and HindII digested pJS208 | This study |

| | | | | | |
|---|---|---|---|---|---|
| pJS208 | Intermediate for pLS80 | Kanamycin, ampicillin | In-fusion ligation | KanR gene and P*aphT* of JC43 and *tetR*-P*tetR* were ligated into EcoRV and KpnI digested pUC18T-mini-Tn7 | This study |

**Table S4: List of synthesized DNA fragments used in this study**

| **name** | **purpose** | **sequence** | **Cloning strategy** |
|---|---|---|---|
| LS4G2 $V_H$ | Generation of pLS06 | CGGCCGCCCACCATGAAATGCAGCTGGGTCATCTTCTTCCTGATGGCAGTGGTTATAGGAATCAATTCAGAGGTTCAGCTGCAGCAGTCTGGGGCAGAGCTTGTGAGGTCAGGGGCCTCAGTCAAGTTGTCCTGCACAGCTTCTGGCTTCAACATTAAAGACTACTATATGGATTGGGTGAAGCAGAGGCCTGAACAGGGCCTGGAGTGGATTGGATGGATTGATCCTGAGAATGGTGATAGTGAATATGCCCCGAAGTTCCAGGGCAAGGCCACTATGACTGCAGACACATCCTCCAACACAGCCTACCTGCAGCTCAGCAGCCTGACATCTGAGGACACTGCCGTCTATTACTGTAATGCCGGACAGCTCGGGCTAGGAGCTTACTGGGGCCAAGGGACTCTGGTCACTGTCTCTGCAGGTGAGTCCTTACAACCTCTTTCTTCTATTCACCTCATTTAGACTTTACTGCATTTGTTGGGGGGGAAATGTGTGTGTCTGAATATGCATGTGCTTACCCTGCCTGTCTCAAAGCAATTGA | Restriction digest using NotI and MfeI |
| LS4G2 $V_L$ | Generation of pLS09 | GCGGCCGCCCACCATGGAATCACAGACCCAGGTCCTCATGTTTCTTCTGCTCTGGGTATCTGGTGCCTGTGCAGACATTGTGATGACACAGTCTCCATCCTCCCTGGCTATGTCAGTAGGACAGAAGGTCACTATGAGCTGCAAGTCCAGTCAGAGCCTTTTAAATAGTAGCAATCAAAAGAACTATTTGGCCTGGTACCAGCAGAAACCAGGACAGTCTCCTAAACTTCTGGTATACTTTGCATCCACTAGGGAATCTGGGGTCCCTGATCGCTTCATAGGCAGTGGATCTGGGACAGATTTCACTCTTACCATCAGCAGTGTGCAGGCTGAAGACCTGGCAGATTACTTCTGTCAGCAACATTATAGCACTCCTCGGACGTTCGGTGGAGGCACCAAGCTGGAAATCAAACGTAAGTATAATCGAATTCGATATCAAGCTT | Restriction digest using NotI and HindIII |
| LS5G11 $V_H$ | Generation of pLS34 | GCGGCCGCCCACCATGAAATGCAGCTGGGTCATCTTCTTCCTGATGGCAGTGGTTATAGGAATCAATTCACAGGTTCAGCTGCAGCAGTCTGGAGCTGAGCTGATGAAGCCTGGGGCCTCAGTGAAGATATCCTGCAAGGCTACTGGCTACAGATTCAGTAGCTACTGGATAGAGTGGGTAAAGCAGAGGCCTGGACAT | Restriction digest using NotI and MfeI |

| | | | |
|---|---|---|---|
| | | GGCCTTGACTGGATTGGAGAGATTTTACCTGGA<br>AGTGGTAGTACTAACTACAATGAGAGGTTCAAG<br>GGCAAGGCCACATTCACTACAGATACATCCTCC<br>AACACAGCCTACATGCAACTCAGCAGCCTGACA<br>TCTGAGGACTCTGCCGTCTATTTCTGTGCAAGA<br>GGAATTTACGGTAGTAGCTTCCTCTTTGACTACT<br>GGGGCCAAGGCACCACTCTCGCAGTCTCCTCAG<br>GTGAGTCCTTACAACCTCTTTCTTCTATTCACCT<br>CATTTAGACTTTACTGCATTTGTTGGGGGGGAAA<br>TGTGTGTGTCTGAATATGCATGTGCTTACCCTGC<br>CTGTCTCAAAGCAATTG | |
| LS5G11 $V_L$ | Generation of pLS35 | GCGGCCGCCCACCATGGAATCACAGACCCAGG<br>TCCTCATGTTTCTTCTGCTCTGGGTATCTGGTGC<br>CTGTGCAGACATTGTGATGTCACAGTCTCCATCC<br>TCCCTACCTGTGTCAGTTGGAGAGAAGGTTACT<br>ATGAGCTGCAAGTCCAGTCAGAGCCTTTTATATA<br>GTAGCAATCAAATGAACTACTTGGCCTGGTACCA<br>GCAGAAACCAGGTCAGTCTCCTAAACTGCTGAT<br>TCACTGGGCATCCACTAGGGAATCTGGGGTCCC<br>TGATCGCTTCACAGGCAGTGGATCTGGGACAGA<br>TTTCACTCTCACCATCAGCAGTGTGAAGGCTGAA<br>GACCTGGCAGTTTATTACTGTCAGCAATATTATA<br>CCTATCCGCTCACGTTCGGTGCTGGGACCAGGC<br>TGGAGCTGAATCGTAAGTATAATCGAATTCGATA<br>TCAAGCTT | Restriction digest using NotI and HindIII |
